# Multivariate GWAS of psychiatric disorders and their cardinal symptoms reveal two dimensions of cross-cutting genetic liabilities

**DOI:** 10.1101/603134

**Authors:** Travis T. Mallard, Richard K. Linnér, Andrew D. Grotzinger, Sandra Sanchez-Roige, Jakob Seidlitz, Aysu Okbay, Ronald de Vlaming, S. Fleur W. Meddens, Bipolar Disorder Working Group of the Psychiatric Genomics Consortium, Abraham A. Palmer, Lea K. Davis, Elliot M. Tucker-Drob, Kenneth S. Kendler, Matthew C. Keller, Philipp D. Koellinger, K. Paige Harden

**Affiliations:** Department of Psychology, University of Texas at Austin, Austin, TX, USA; Department of Economics, School of Business and Economics, Vrije Universiteit Amsterdam, Amsterdam, The Netherlands; Autism and Developmental Medicine Institute, Geisinger, Lewisburg, PA, USA; Department of Psychiatry, University of California San Diego, La Jolla, CA, USA; Department of Child and Adolescent Psychiatry and Behavioral Science, Children’s Hospital of Philadelphia, Philadelphia, PA, USA; Department of Psychiatry, University of Pennsylvania, Philadelphia, PA, USA; Erasmus University Rotterdam Institute for Behavior and Biology, Erasmus School of Economics, Erasmus University Rotterdam, Rotterdam, The Netherlands; Institute for Genomic Medicine, University of California San Diego, La Jolla, CA, USA; Division of Genetic Medicine, Department of Medicine, Vanderbilt University Medical Center, Nashville, TN, USA; Vanderbilt Genetics Institute, Vanderbilt University Medical Center, Nashville, TN, USA; Population Research Center, University of Texas at Austin, Austin, TX USA; Virginia Institute for Psychiatric and Behavioral Genetics, Virginia Commonwealth University, Richmond, VA, USA; Department of Psychiatry, Medical College of Virginia/Virginia Commonwealth University, Richmond, VA, USA; Institute for Behavioral Genetics, University of Colorado Boulder, Boulder, CO, USA; Department of Psychology and Neuroscience, University of Colorado Boulder, Boulder, CO, USA

## Abstract

Understanding which biological pathways are specific versus general across diagnostic categories and levels of symptom severity is critical to improving nosology and treatment of psychopathology. Here, we combine transdiagnostic and dimensional approaches to genetic discovery for the first time, conducting a novel multivariate genome-wide association study (GWAS) of eight psychiatric symptoms and disorders broadly related to mood disturbance and psychosis. We identify two transdiagnostic genetic liabilities that distinguish between common forms of mood disturbance (major depressive disorder, bipolar II, and self-reported symptoms of depression, mania, and psychosis) versus rarer forms of serious mental illness (bipolar I, schizoaffective disorder, and schizophrenia). Biological annotation revealed divergent genetic architectures that differentially implicated prenatal neurodevelopment and neuronal function and regulation. These findings inform psychiatric nosology and biological models of psychopathology, as they suggest the severity of mood and psychotic symptoms present in serious mental illness may reflect a difference in kind, rather than merely in degree.

---

Psychiatric disorders are one of the leading causes of global disease burden, affecting more than 25% of the world’s population at some point during their lifetime^1^. Twin- and family-based studies have established that a substantial portion of individual differences in liability to psychiatric disorders is caused by genetic variation^2^. Genome-wide association studies (GWASs) have identified numerous genetic loci that have replicable associations with severe and debilitating psychiatric disorders, including schizophrenia^3^, bipolar disorder^4^, and major depressive disorder^5^.

GWASs have also identified a substantial degree of genetic overlap across psychiatric disorders, finding high genetic covariances and many pleiotropic loci^6,7^. This genetic overlap complicates efforts to identify causes, consequences, and treatments that are specific to any individual psychiatric disorder^8^. In response to these challenges, transdiagnostic approaches to psychiatric disease aim to identify biological systems that are perturbed across many forms of illness^9,10^. Transdiagnostic research may yield new therapeutic targets with broad utility, as well as inform nosological classification and stratification of at-risk populations.

Concurrent with the emergence of transdiagnostic research, efforts to identify disorder-specific genetic loci have turned toward studying self-report measures in the general population^11–13^, as case-control study designs require diagnostic schedules that can be slow and costly. If valid, this dimensional approach in non-clinical samples has the potential to accelerate genetic discovery via dramatic increases in sample size, as self-report survey measures of psychiatric symptoms can be administered at-scale to large, genotyped population-based samples, such as UK Biobank^14,15^. However, while this approach may be valid for some common forms of psychopathology^16^, it is unknown whether the biology that influences normative variation in subthreshold symptoms also underlies rarer psychiatric conditions, such as those characterized by mania and/or psychosis.

Here, we combine transdiagnostic and dimensional research approaches to genetic discovery by analyzing the genetic relationships among eight psychiatric phenotypes related to mood and psychotic disorders: depressive symptoms, manic symptoms, psychotic symptoms, major depressive disorder, bipolar II disorder, bipolar I disorder, schizoaffective disorder, and schizophrenia. In doing so, we aim to address three main questions. First, what is the genetic basis of mood and psychotic symptoms in the general population and how does it compare to the genetic basis of psychiatric diagnoses that are characterized by those symptoms? Second, how many transdiagnostic dimensions of genetic liability cut across these eight phenotypes? Third, how are dimensions of transdiagnostic liability similar and dissimilar in their genetic architecture, underlying biology, and associations with other aspects of human well-being and disease?

## Results

### Novel loci associated with lifetime endorsement of mood and psychotic symptoms

We used a combination of Bayesian item response theory and linear mixed models to conduct univariate GWAS for self-reported measures of lifetime depression, mania, and psychosis from 252,252 individuals in the UK Biobank. We observed substantial inflation of the median test statistic for all three phenotypes, and the linkage disequilibrium (LD) score regression intercepts and attenuation ratios suggest that test statistic inflation is primarily due to polygenic signal rather than bias (Figure 1, Table 1). After applying a standard clumping algorithm via FUMA (*r*^2^ = .1, 250kb merge window), we identified 23 independent loci associated with lifetime depressive, manic, and/or psychotic symptoms (Supplementary Tables 1-3). Nine of these loci were significantly associated with two or more phenotypes, and six loci were associated with all three psychiatric phenotypes.

**Table 1.** Summary of study phenotypes.

| **Univariate GWAS (abbr.)** | **Source** | ***N*** | **$h^2$** | **$\lambda_{GC}$** | **Mean $\chi^2$** | **Intercept** | **Ratio** |
| --- | --- | --- | --- | --- | --- | --- | --- |
| Depressive symptoms (DEP) | Present study | 252,252 | .08 | 1.31 | 1.38 | 1.01 | .02 |
| Manic symptoms (MAN) | Present study | 252,252 | .08 | 1.31 | 1.39 | 1.00 | .00 |
| Psychotic symptoms (PSY) | Present study | 252,252 | .07 | 1.31 | 1.33 | 1.00 | .01 |
| Major depressive D/O (MDD) | Wray *et al.*, 2018<sup>5</sup> | 138,884 | .10 | 1.19 | 1.20 | 1.00 | < 0 |
| Bipolar II D/O (BD2) | Stahl *et al.*, 2019<sup>4</sup> | 25,576 | .10 | 1.07 | 1.08 | 1.03 | .42 |
| Bipolar I D/O (BD1) | Stahl *et al.*, 2019<sup>4</sup> | 45,871 | .22 | 1.31 | 1.37 | 1.04 | .09 |
| Schizoaffective D/O (SZA) | Stahl *et al.*, 2019<sup>4</sup> | 9,667 | .27 | 1.06 | 1.06 | 1.02 | .35 |
| Schizophrenia (SCZ) | Ruderfer *et al.*, 2018<sup>20</sup> | 65,967 | .23 | 1.49 | 1.63 | 1.05 | .08 |
| **Multivariate GWAS** | **Source** | **$N_{\text{eff}}$** |  | **$\lambda_{GC}$** | **Mean $\chi^2$** | **Intercept** | **Ratio** |
| F1 (Mood Disturbance) | Present study | 377,518 |  | 1.44 | 1.53 | 1.05 | .10 |
| F2 (Serious Mental Illness) | Present study | 51,276 |  | 1.46 | 1.61 | 1.02 | .04 |

**Note:** Heritability ( $h^2$ ) was estimated using LD score regression and is reported on the observed scale. $\lambda_{GC}$ refers to the median $\chi^2$ statistic of the GWAS divided by the expected median of the $\chi^2$ distribution with 1 degree of freedom. Mean $\chi^2$ refers to the average $\chi^2$ statistic of the GWAS. Intercept refers to the estimated intercept from univariate LD score regression. Ratio refers to a measure of stratification bias that is defined as (Intercept – 1) / (Mean $\chi^2$ – 1). To harmonize measurement approaches among psychiatric disorders, summary statistics for MDD were obtained for the clinically-ascertained cohorts, excluding 23andMe and UK Biobank. D/O = disorder. $N_{\text{eff}}$ refers to effective sample size (Method).

**Figure 1.**
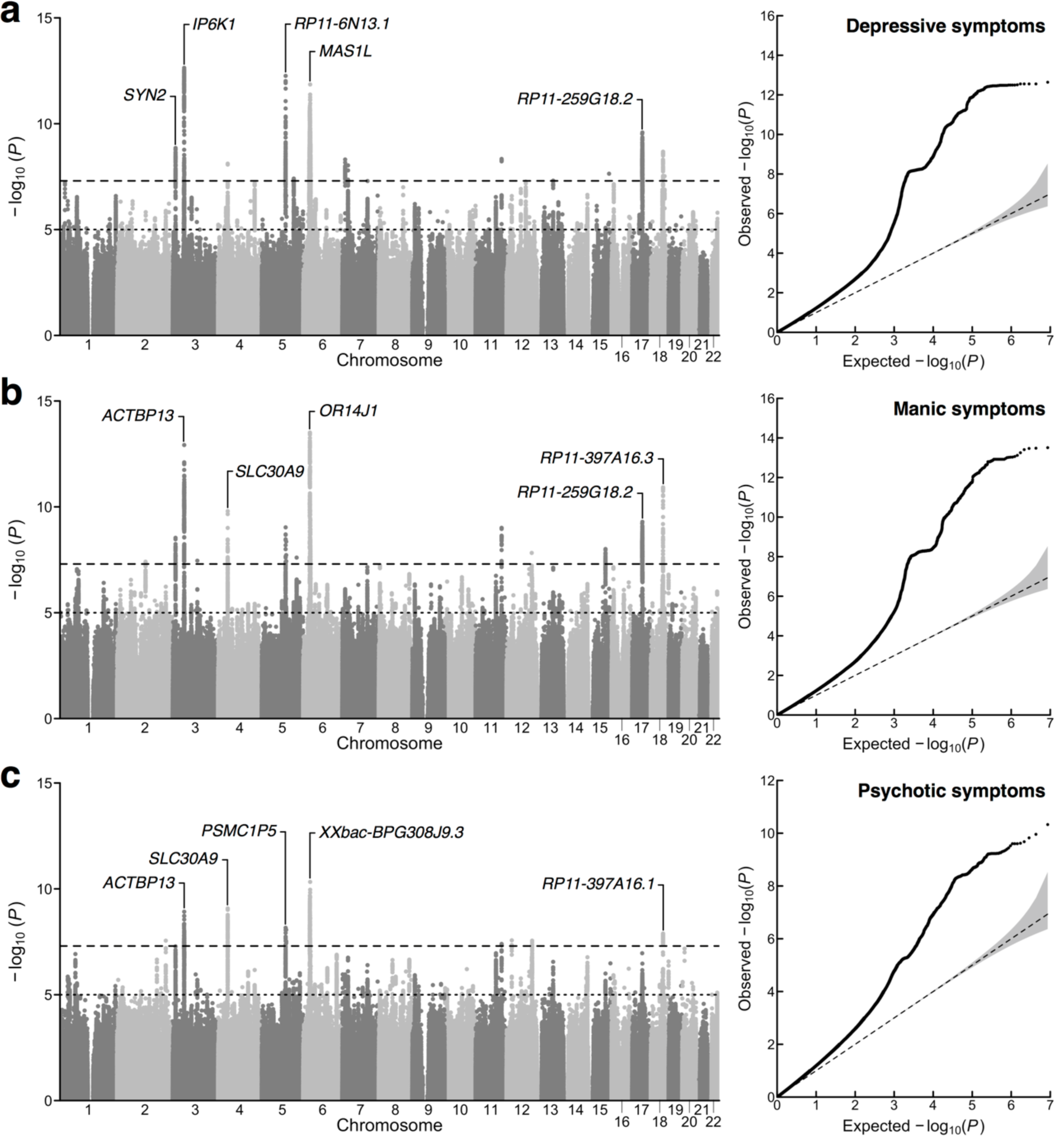
Univariate association results for lifetime measures of mood disturbance and psychosis. **a**,**b**,**c**, Manhattan plots and a quantile-quantile plots for (**a**) depressive, (**b**) manic, and (**c**) psychotic symptoms. In the Manhattan plots, the x-axis refers to chromosomal position, the y-axis refers to the significance on a -log_10_ scale, the horizontal dashed line denotes genome-wide significance (*P* = 5e-8), and the horizontal dotted line marks suggestive significance (*P* = 1e-5). In the quantile-quantile plots, the x-axis refers to expected *P* value, while the y-axis refers to the observed *P* value. For each plot, the nearest gene for the lead SNP in the top five genome-wide significant loci is labeled.

The identified risk loci span across 12 chromosomes and include variants tagging the major histocompatibility complex region on chromosome 6 and a well-known inversion polymorphism on chromosome 17 previously associated with several psychiatric phenotypes^17^. Many of these risk loci replicated previous findings from GWASs of psychopathology or were in high LD with previous hits for phenotypes including neuroticism^18^ (*e.g*., rs7111031, rs10503002, rs4245154), broadly defined depression^11^ (*e.g*., rs9586, rs191800971, rs7111031), and schizophrenia^19^ (*e.g*., rs1233494, rs4245154, rs4702). However, several loci contained lead SNPs that were new GWAS signals altogether, identifying new regions of the genome that confer risk for psychopathology, such as rs4722389, rs7324564, and rs570217967.

Moreover, our gene-based association analyses performed via MAGMA identified 144 genes associated with at least one of the psychiatric symptoms (depression, mania, or psychosis), 39 of which were associated with all three. For all phenotypes, we observed enriched expression in brain tissue, as well as an enriched signal for brain-related gene sets. We report detailed biological annotation (*e.g*., gene mapping, gene set enrichment, tissue enrichment) for each of these GWASs in Supplementary Tables 4-12.

### Two transdiagnostic genetic liabilities underlie mood and psychotic psychopathology

To characterize the genomic relationships among psychiatric symptoms and disorders commonly characterized by depression, mania, and/or psychosis^4,5,20^ (see Table 1 for overview of study phenotypes), we first used bivariate LD score regression to estimate genetic correlations between all pairs of psychiatric phenotypes. We observed very large positive genetic correlations among the three psychiatric symptoms (mean *r*_g_ = .95, *SEM* = .02); however, we observed more modest genetic correlations for the five psychiatric disorders (mean *r*_g_ = .55, *SEM* = .09). We found that schizophrenia, schizoaffective disorder, and bipolar I were highly correlated with one another (*r*_gSCZ-SZA_ = .87 [*SE* = .13], *r*_gSCZ-BD1_ = .72 [*SE* = .03], *r*_gSZA-BD1_ = .81 [*SE* = .12]), but these disorders generally had markedly smaller genetic correlations with bipolar II and major depressive disorder (*r*_gSCZ-BD2_ = .53 [*SE* = .03], *r*_gSCZ-MDD_ = .39 [*SE* = .04], *r*_gSZA-BD2_ = .28 [*SE* = .21], *r*_gSZA-MDD_ = .06 [*SE* = .12], *r*_gBD1-MDD_ = .33 [*SE* = .04]); however, bipolar I and bipolar II were highly correlated (*r*_gBD1-BD2_ = .88 [*SE* = .11]). Interestingly, we found that bipolar II and major depressive disorder were highly correlated with each other (*r*_gBD2-MDD_ = .69 [*SE* = .13]), as well as with all psychiatric symptoms (*r*_gBD2-DEP_ = .75 [*SE* = .11], *r*_gBD2-MAN_ = .71 [*SE* = .11], *r*_gBD2-PSY_ = .70 [*SE* = .11], *r*_gMDD-DEP_ = .85 [*SE* = .03], *r*_gMDD-MAN_ = .77 [*SE* = .03], *r*_gMDD-PSY_ = .80 [*SE* = .04]).

After applying a hierarchical clustering algorithm to the genetic correlation matrix, we found two distinct clusters of psychiatric phenotypes (Figure 2a). The first cluster comprised the three psychiatric symptoms, major depressive disorder, and bipolar II, and the second cluster comprised bipolar I, schizoaffective disorder, and schizophrenia. We then conducted an exploratory factor analysis (EFA) of the genetic covariance matrix. EFA results were consistent with the groupings suggested by the hierarchical clustering algorithm, identifying a correlated two-factor model with approximate simple structure. That is, we found that phenotypes principally loaded onto one of two latent genetic factors with negligible cross-loadings (Figure 2b). Combined, these two correlated latent factors explained 81.3% of the total genetic variance across phenotypes. Finally, we formally modeled the genetic covariance matrix via confirmatory factor analysis (CFA). We based our model on the EFA results, which consisted of two correlated latent factors, F1 and F2. F1 can be conceptualized as capturing common psychopathology related to mood disturbance (including self-reported depressive, psychotic, and manic symptoms, as well as bipolar II and major depressive disorder), while F2 can be conceptualized as capturing rarer forms of serious mental illness (bipolar I, schizoaffective disorder, and schizophrenia). We did not estimate any cross-loadings. Instead, we estimated correlated residuals between bipolar I and bipolar II, as inspection of the genetic correlation matrix suggested a unique relationship between these disorders. The path diagram for this model is presented in Figure 2c.

**Figure 2.**
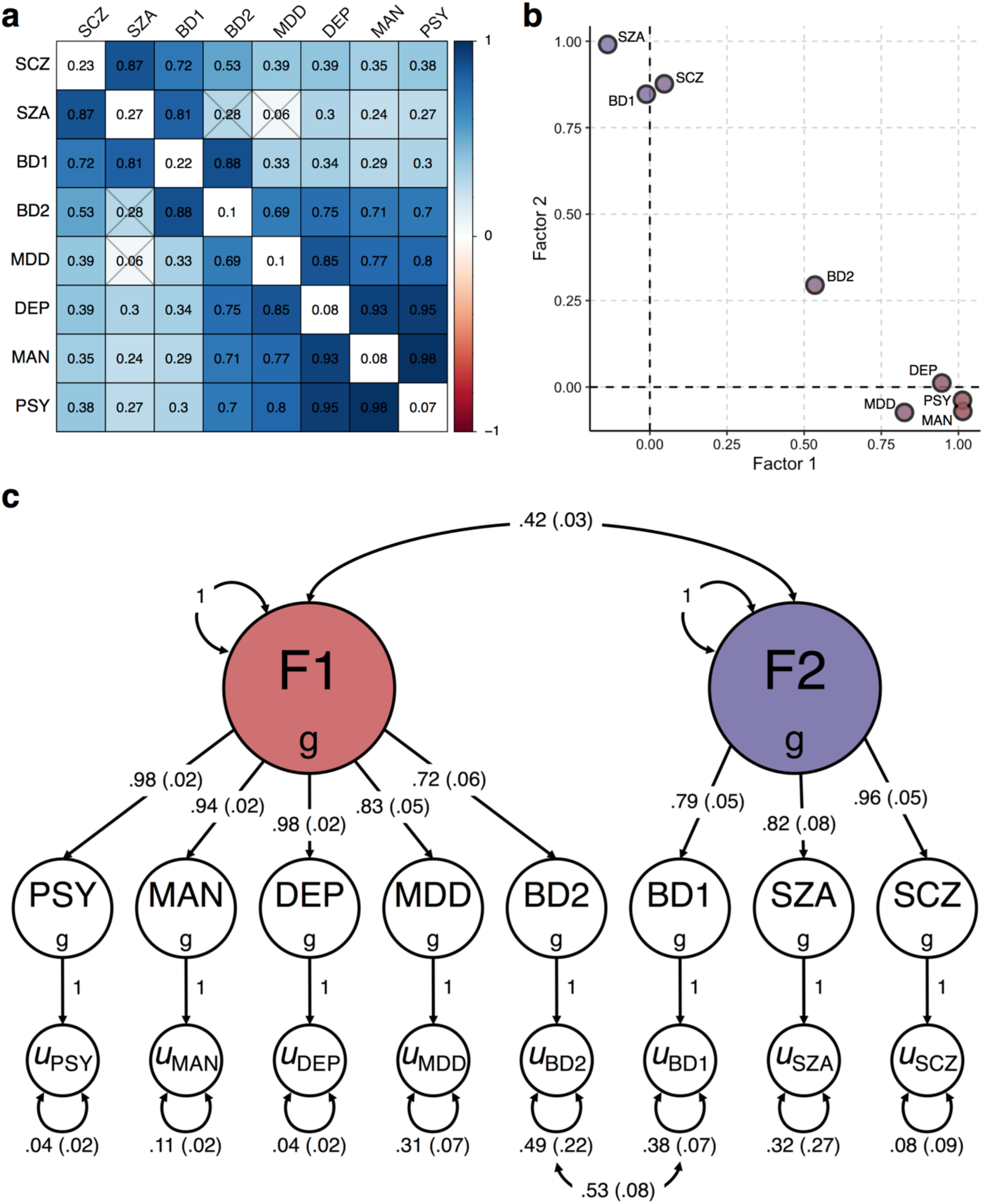
Relationships between eight psychiatric symptoms and disorders. **a**, Matrix of bivariate genetic correlation estimates, where the diagonal elements correspond to SNP *h*^2^ and the off-diagonal elements correspond to genetic correlations. Estimates that are non-significant are crossed out. **b**, Scatterplot of standardized factor loadings from the exploratory factor analysis. **c**, Path diagram for the final confirmatory factor model with standardized parameter estimates.

We compared the correlated factor model to a common factor model, where all phenotypes are indicators of a single latent factor (*i.e*., a *p* factor) (Supplementary Section 1.1). Briefly, we found that the common factor model showed suboptimal fit to the data, while the correlated factors model with correlated residuals for bipolar I and bipolar II showed excellent fit. Fit indices from the CFA indicated that the correlated factors model closely approximated the observed genetic covariance matrix (χ2(18) = 496.16, AIC = 532.16, CFI = .99, SRMR = .06). That is, the patterns of covariance among the eight psychiatric phenotypes were most parsimoniously represented by two transdiagnostic latent factors at the genetic level, which were correlated only modestly (*r*_g_ = .42; *SE* = .03). This is a notable divergence from the factor structure frequently observed at the phenotypic level, including that seen with similar phenotypes in the UK Biobank (Supplementary Section 1.1, Supplementary Figures 1-3).

### Transdiagnostic factors have markedly divergent genetic architectures

We then conducted a multivariate GWAS that estimated SNP associations with the latent factors, F1 (*N*_*eff*_ = 377,518) and F2 (*N*_*eff*_ = 51,276). Multivariate GWAS results are presented in Table 1 and Figure 3. We observed substantial inflation of the median test statistic for both F1 (λ_GC_ = 1.44, mean χ^2^ = 1.53) and F2 (λ_GC_ = 1.46, mean χ^2^ = 1.61), indicative of a robust polygenic signal for both factors (Supplementary Figure 4). The LD Score regression intercepts and attenuation ratios for F1 (intercept = 1.05, *SE* = .01; ratio = .10, *SE* = .02) and F2 (intercept = 1.02, *SE* = .01; ratio = .04, *SE* = .02) suggest that test statistic inflation is primarily due to polygenic signal rather than bias.

**Figure 3.**
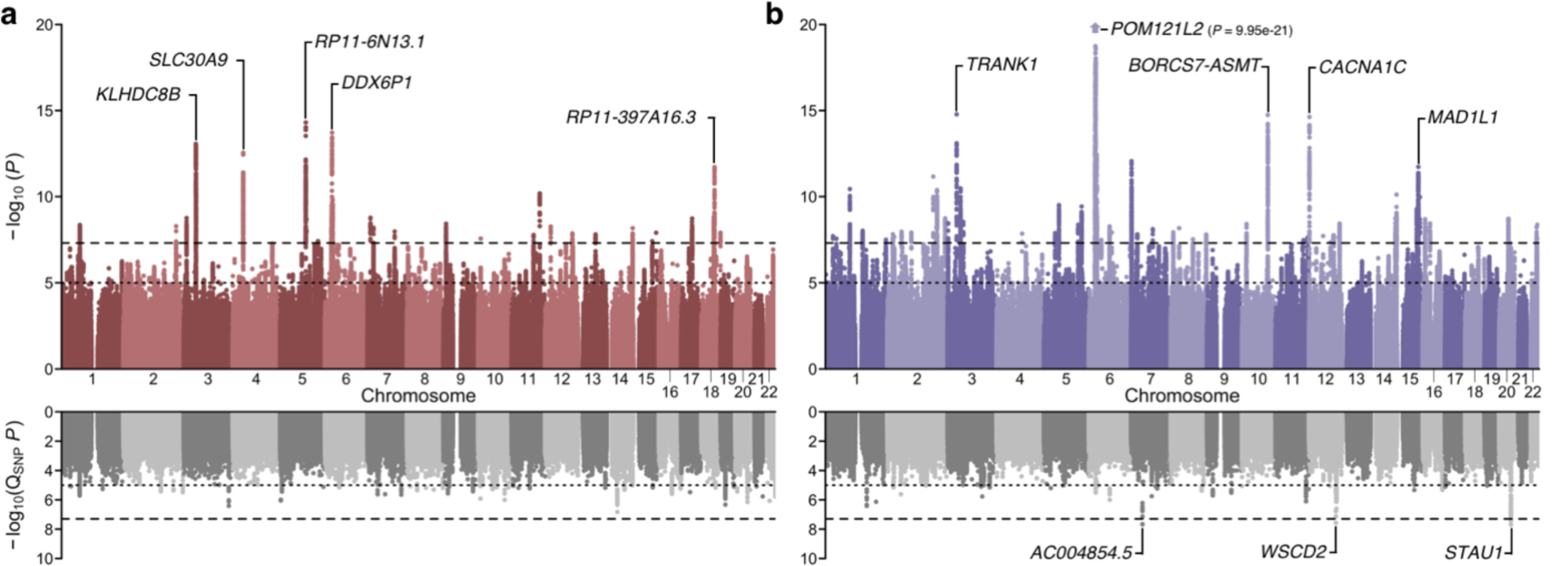
Multivariate association results for the two transdiagnostic latent genetic factors. **a**,**b**, Miami plots for (**a**) F1 and (**b**) F2. The top of each Miami plot corresponds to the significance of SNP effects on each latent factor, as traditionally conveyed in a Manhattan plot, while the bottom corresponds to the significance of heterogeneity tests for SNP effects (Q_SNP_; *i.e*., the degree to which SNP effects are not mediated by F1 or F2). For each plot, the x-axis refers to chromosomal position, the y-axis refers to the significance on a -log_10_ scale, the horizontal dashed line denotes genome-wide significance (*P* = 5e-8), and the horizontal dotted line marks suggestive significance (*P* = 1e-5). For each plot, the nearest gene for the lead SNP in the top five genome-wide significant loci is labeled.

**Figure 4.**
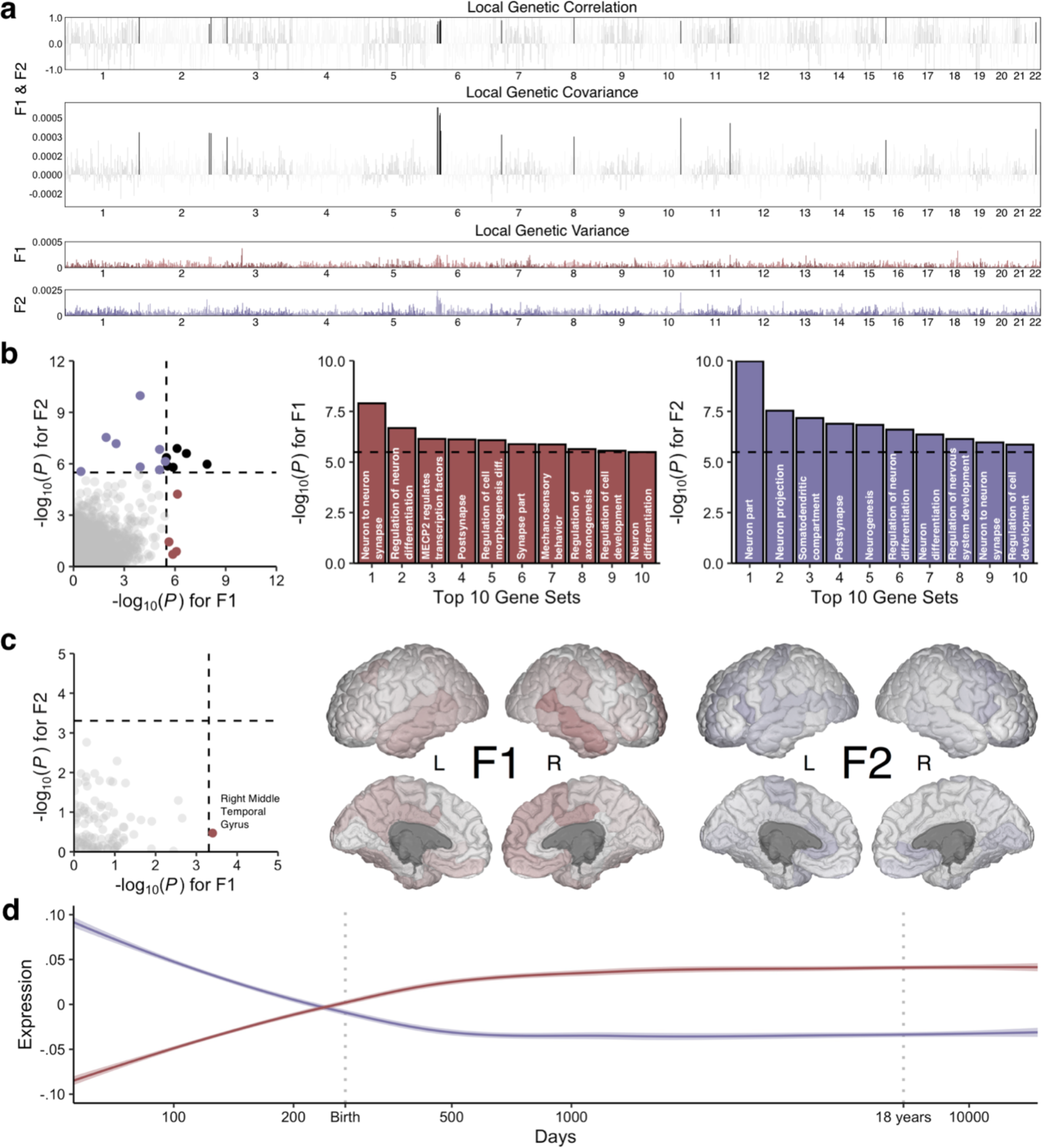
Biological annotation of the two transdiagnostic latent genetic factors. **a**, Manhattan plots for local genetic correlation, covariance, and variance for F1 and F2. Black bars indicate significant local genetic correlation. **b**, Scatterplot of gene set enrichment results illustrating convergence and divergence across the latent genetic factors with accompanying histograms for the top 10 gene sets for each factor. **c**, Scatterplot of neuroimaging genetic correlation results with accompanying figures where the -log_10_ *P* values are mapped across the cortex, as parcellated in the Desikan-Killiany-Tourville atlas. **d**, Smoothed line plots of gene set expression across developmental time in the Brainspan dataset for prioritized genes with transcriptomic profiles that are spatially similar to the neuroimaging genetic correlation maps for F1 and F2 (as indexed in the Allen Human Brain Atlas). For all plots, the dashed black line corresponds to the Bonferroni-corrected significance threshold when applicable.

We applied a standard clumping algorithm and identified 26 and 59 independent loci associated with F1 and F2, respectively (Table 2, Supplementary Tables 13-14). Only 5 loci were associated with both F1 and F2. While many of these genomic regions have been previously identified in either the constituent GWASs or related studies, several contain novel discoveries. For example, 4 of the 26 loci associated with F1 contain lead SNPs that have not been previously associated with psychopathology: rs13153844 (*P* = 2.09e-9, nearest gene = *PSMC1P5*), rs1551765 (*P* = 3.89e-8, nearest gene = *GRIA1*), rs147584788 (*P* = 1.08e-8, nearest gene = *AC003088.1*), rs8035987 (*P* = 3.94e-8, nearest gene = *SIN3A*). Similarly, 10 of the 59 loci associated with F2 contain lead SNPs that are also novel risk variants for psychopathology: rs2953329 (*P* = 3.27e-8, nearest gene = *AKT3*), rs10199182 (*P* = 1.56e-8, nearest gene = *AC068490.2*), rs9463650 (*P* = 3.34e-8, nearest gene = *RPS17P5*), rs12190758 (*P* = 1.22e-8, nearest gene = *RP1-149C7.1*), rs13233308 (*P* = 9.20e-9, nearest gene = *ABCB1*), rs11603014 (*P* = 2.32e-8, nearest gene = *RP11-890B15.2*), rs11104379 (*P* = 4.50e-8, nearest gene = *RPL23AP68*), rs10777957 (*P* = 1.79e-8, nearest gene = *ANKS1B*), rs11064837 (*P* = 2.43e-8, nearest gene = *RP11-768F21.1*), and rs11908600 (*P* = 4.97e-8, nearest gene = *STK4*).

**Table 2.**
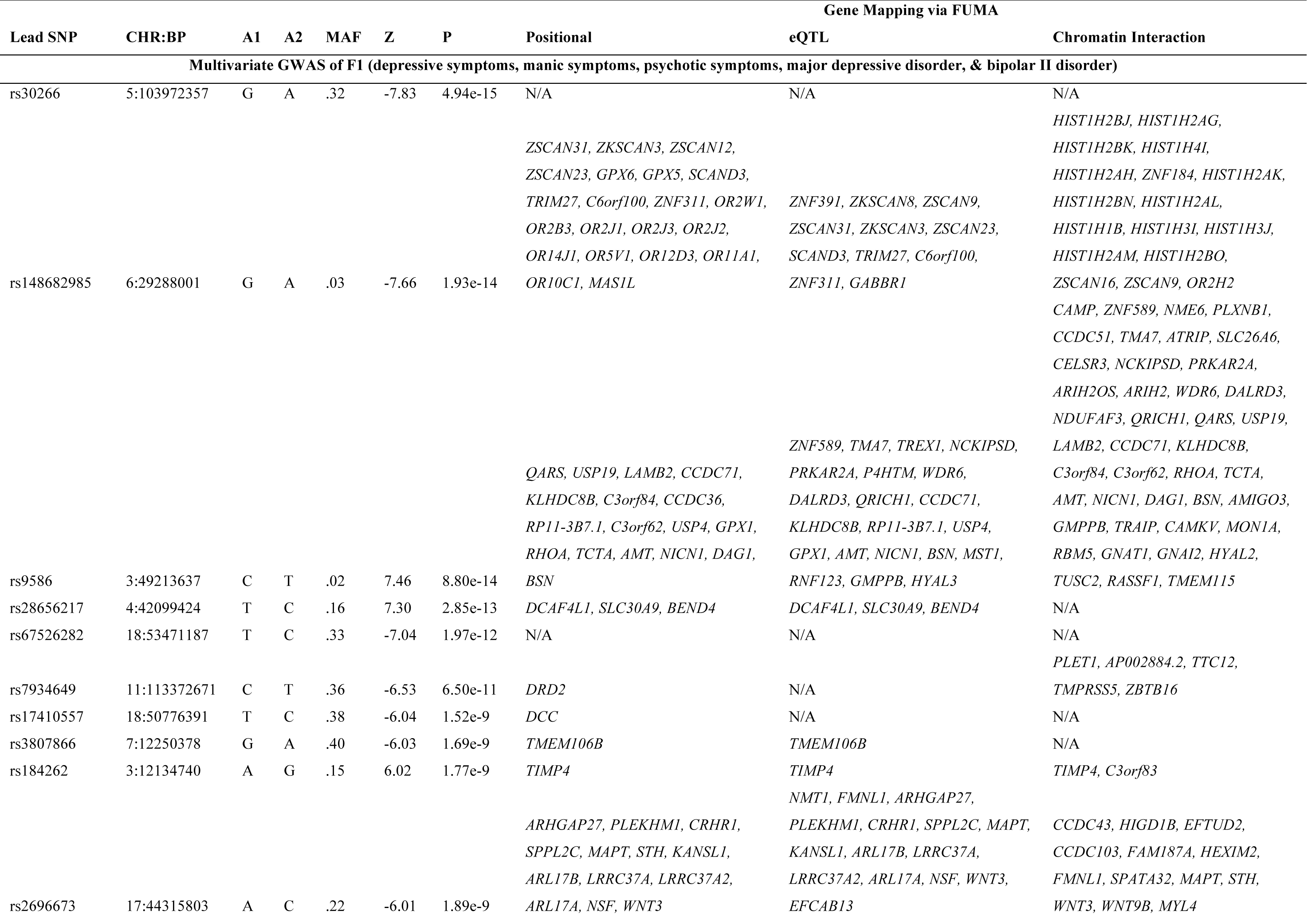
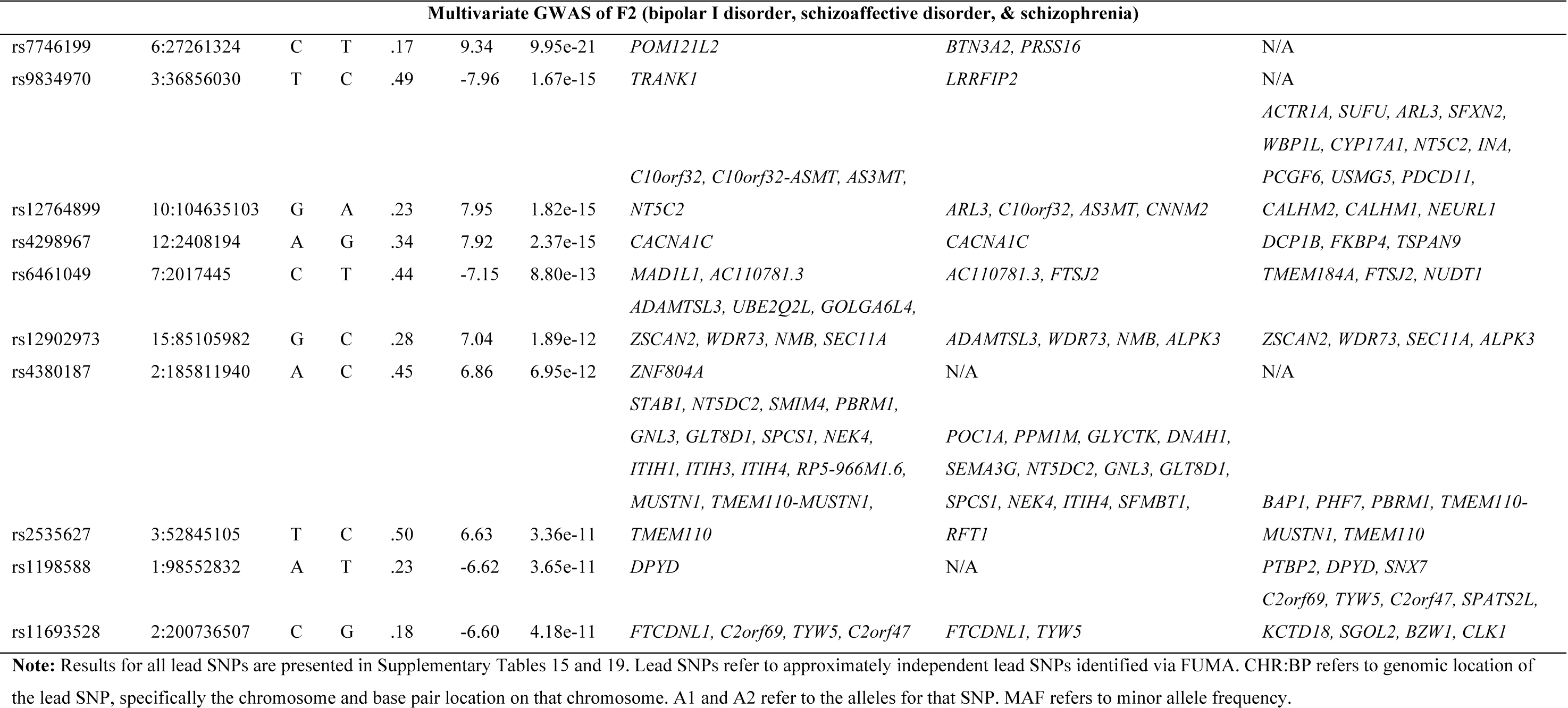
Lead SNPs for the top ten loci per latent factor from multivariate association analyses.

Tests of heterogeneity suggested that the majority of observed SNP effects operate via the latent factors (*i.e*., associated SNPs primarily had consistent, pleiotropic effects on the constituent phenotypes). Indeed, *Q*_SNP_ tests identified no heterogeneous loci for F1 and only three heterogeneous loci for F2 with lead SNPs rs11696888 on chromosome 20 (*Q*_SNP_ *P* = 2.04e-8; nearest gene = *STAU1*), rs1990042 on chromosome 7 (*Q*_SNP_ *P* = 2.10e-8, nearest gene = *AC004854.1*), and rs3764002 on chromosome 12 (*Q*_SNP_ *P* = 2.70e-8; nearest gene = *WSCD2*). Interestingly, the heterogeneous locus with lead SNP rs11696888 also contains rs200005157, which is a four base-pair insertion/deletion that was previously identified as a locus with divergent effects on bipolar disorder and schizophrenia^20^. Fine-mapping conducted by Ruderfer and colleagues identified *CSE1L* as a plausible causal gene with divergent effects for bipolar disorder and schizophrenia on chromosome 20.

### Transdiagnostic factors are related to different aspects of neurobiology

To characterize the effects of variants associated with the transdiagnostic factors of psychopathology, we used FUMA to conduct a series of gene mapping analyses. Specifically, we used positional mapping to align SNPs to genes based on genomic location, expression quantitative trait loci (eQTL) mapping to match cis-eQTL SNPs to genes whose expression they affect, and chromatin interaction mapping to link SNPs to genes on the basis of three-dimensional DNA-DNA interactions. These three methods linked the associated SNPs for F1 and F2 to a combined 287 and 570 putative risk genes, respectively (Supplementary Tables 15-22). We also used MAGMA to conduct gene-based association analyses, which identified 131 and 284 genes associated with F1 and F2 (Supplementary Tables 23-24). Finally, we used S-MultiXcan to identify 50 and 91 genes associated with differential expression levels in brain tissue for F1 and F2, respectively (Supplementary Tables 25-26). Collectively, these five approaches link a total of 344 putative risk genes to F1 and 748 putative risk genes to F2.

When considered as a set, biological annotation of these genes linked genetic risk for psychopathology to the central nervous system. Briefly, we found that mapped genes for F1 and F2 were both linked to brain-associated eQTLs, enriched for gene sets broadly related to regulatory biological processes, and previously identified in myriad GWAS related to psychopathology, cognition, and brain morphology and health (see Supplementary Tables 27-28). These results provide preliminary evidence for how risk variants for both genetic liabilities are functionally related to the brain and related neuropsychiatric phenotypes. Perhaps more importantly, they further highlight the relatively modest overlap in shared genetic architecture, as only 17% (155/937) of the unique putative risk genes were linked to both F1 and F2.

As previously noted, F1 and F2 were only modestly correlated with one another (*r*_g_ = .42, *SE* = .03) implying that the majority of genetic variance in each factor is unique from the other. To further characterize the shared and unique genetic architecture of F1 and F2, we used HESS to estimate the local genetic covariance for 1,698 contiguous, similarly-sized partitions across the genome. We found that approximately 27% of the genome explains 80% of the total genetic covariance between F1 and F2, and only 15 genomic partitions share a significant local genetic correlation after correcting for multiple comparisons (Figure 4a). Collectively, these results further emphasize that these latent genetic factors are largely distinct from one another.

Gene-set enrichment and gene property (*i.e*., tissue expression) analyses further suggest that the genetic architectures of F1 and F2 are divergent at more granular levels of analysis, converging only at higher levels. While results from gene set enrichment analyses broadly implicated neurodevelopmental and neurobiological pathways for both factors, the specific molecular functions, cellular components, and biological processes were generally different (Figure 4b, Supplementary Tables 29-30). For example, gene sets related to neurons were enriched for F1 and F2, but gene sets for specific parts of neurons were differentially enriched (*e.g*., the axon for F1 versus the somatodendritic compartment for F2). Similarly, in the tissue expression analysis, we found that the brain was broadly implicated in the pathogenesis of psychopathology, as nearly all brain-related tissues were enriched for both F1 and F2 (Supplementary Tables 31-32). At the level of brain tissue, the only regions with divergent effects were the substantia nigra and brainstem, which were not significantly enriched for F1 after correction for multiple comparisons. However, shortcomings of these analyses include the relatively low spatial resolution of brain-related gene expression data, and the limited sample size of the underlying data.

To gain greater insight into potential etiological relationships between psychopathology and neurobiology, we estimated genetic correlations between the transdiagnostic factors of psychopathology and 101 morphological features of the human brain. Although we generally observed negative genetic correlations with cortical and subcortical features (*i.e*., greater risk for psychopathology is associated with smaller volumes across the brain), and positive with ventricular features (*i.e*., greater risk for psychopathology is associated with larger ventricular volumes), specific estimates between morphological features and F1 and F2 showed relatively little concordance (Figure 4c). After correcting for multiple comparisons, only the genetic correlation between F1 and the right middle temporal gyrus remained statistically significant (*r*_g_ = -.15, *SE* = .04, *P* = 3.98e-4) (Supplementary Table 33).

We then used data from the Allen Human Brain Atlas to identify genes with transcriptomic profiles that were spatially similar to the neuroimaging genetic correlation maps for F1 and F2 (Supplementary Tables 35-36). Notably, these transcriptomically-prioritized gene sets for F1 and F2 were entirely disjoint from one another, and differentially expressed in pre-and postnatal cortical tissue from the Brainspan dataset (Figure 4d). We found that the developmental expression profile of the F1 gene set most closely resembled that of postnatal inhibitory neuronal genes, while the developmental expression profile of the F2 gene set most closely resembled that of prenatal inhibitory neuronal genes^21,22^ (Supplementary Figure 5).

**Figure 5.**
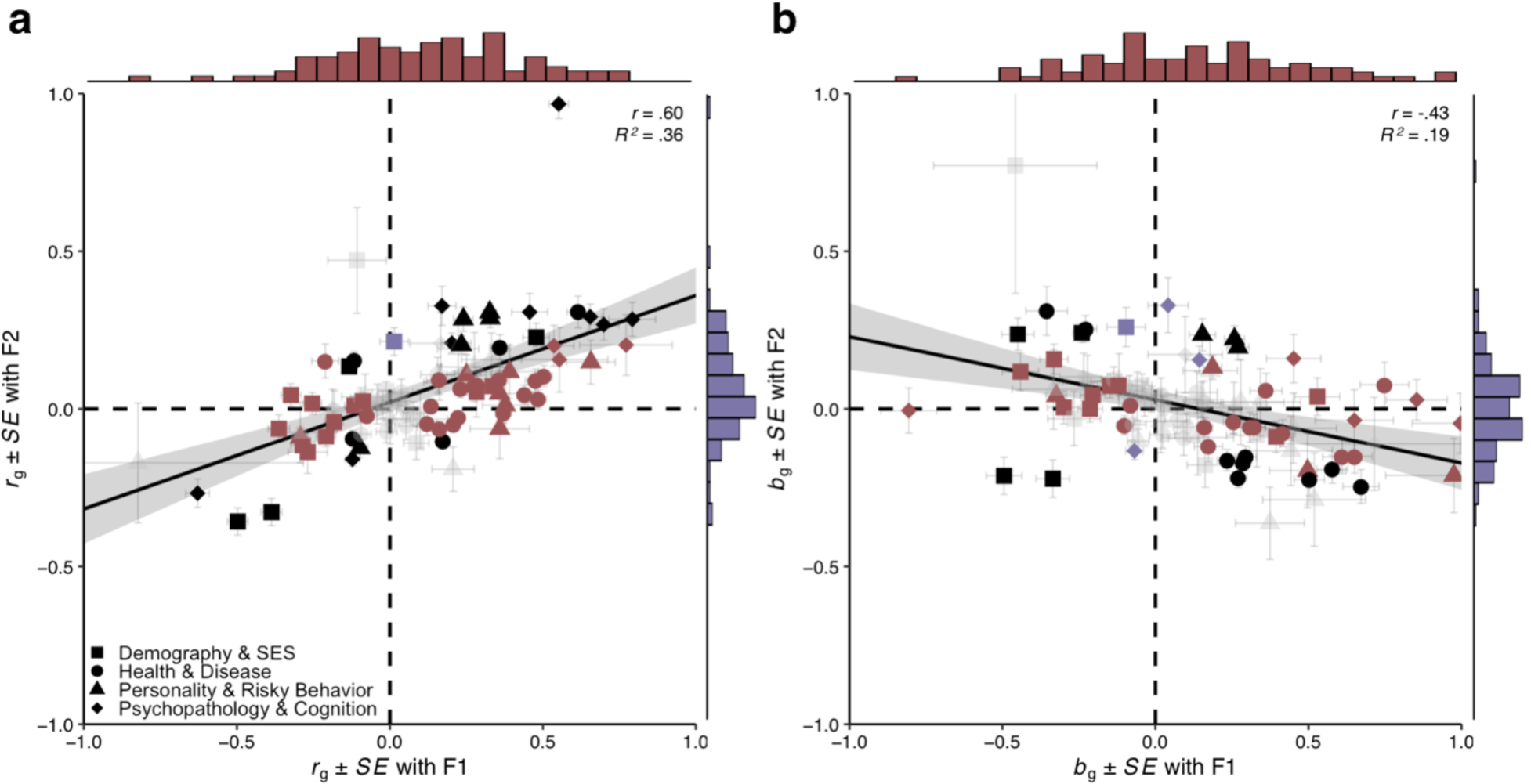
Genetic correlation results for the two transdiagnostic latent genetic factors. **a**, Scatterplot of genetic correlations (*r*_g_) with marginal histograms. **b**, Scatterplot of partial genetic correlations (*b*_g_) with marginal histograms. For both plots, phenotypes are grouped into one of four broad domains: (i) demography and socioeconomic status, (ii) health and disease, (iii) personality and risky behavior, and (iv) psychopathology and cognition. A line-of-best fit (with 95% confidence interval) is fit for all 92 data points. Points are colored burgundy if significant only for F1, violet if significant only for F2, black if significant for both, and faded gray if non-significant for both. The standard errors (*SE*) for point estimates are plotted for both factors.

### Transdiagnostic factors are differentially associated with human health and well-being

To better understand how these transdiagnostic genetic liabilities may manifest above and beyond their constituent phenotypes, we conducted a series of genetic correlation and polygenic prediction analyses focused on theoretically relevant phenotypes. In the genetic correlation analyses, we evaluated the relationships between the latent factors of psychopathology and 92 phenotypes broadly related to four broad domains of human health and well-being (Figure 5a) (Supplementary Table 37). We found that genetic correlation estimates for F1 and F2 were moderately correlated across all broad domains (*r* = .60, *P* = 2.77e-10), as well as within each of the four domains: demography and socioeconomic status (*r* = .55, *P* = 1.17e-2), health and disease (*r* = .42, *P* = 1.17e-2), personality and risky behavior (*r* = .60, *P* = 3.05e-3), and psychopathology and cognition (*r* = .63, *P* = 1.57e-2). Generally, we found that F1 was more consistently correlated with phenotypes typically related to psychopathology than F2. This pattern was also observed in the partial genetic correlation analyses, where we found strong evidence of divergent genetic correlations after accounting for the overlap between F1 and F2 (Figure 5b) (Supplementary Table 38). Indeed, partial genetic correlation estimates for F1 and F2 were negatively correlated across all domains (*r* = -.43, *P* = 2.72e-5).

In the polygenic prediction analyses, we used electronic health records from the Vanderbilt University Medical Center biobank (BioVU) to evaluate the penetrance and pleiotropy of genetic risk for the transdiagnostic factors of psychopathology across 1,335 disease phenotypes, hereby referred to as “phecodes” (Figure 6) (Supplementary Tables 39-40). We found that polygenic scores for F1 and F2 were generally associated with all of the constituent phenotypes for both factors, but F1 was more strongly associated with mood-related phecodes while F2 was more strongly associated with psychosis-related phecodes. Both polygenic scores for F1 and F2 shared associations with some forms of psychopathology (*e.g*., suicidality, posttraumatic stress disorder, substance use disorders, and anxiety disorders), but diverged in their associations with others (*e.g*., personality disorders, paranoid disorders). Beyond psychopathology, F1 was more consistently associated a variety of medical phecodes, including those related to infectious diseases (*e.g*., viral hepatitis, human immunodeficiency virus disease) and pervasive developmental disorders, as well as diseases of the circulatory, digestive, endocrine, genitourinary, musculoskeletal, and respiratory systems.

**Figure 6.**
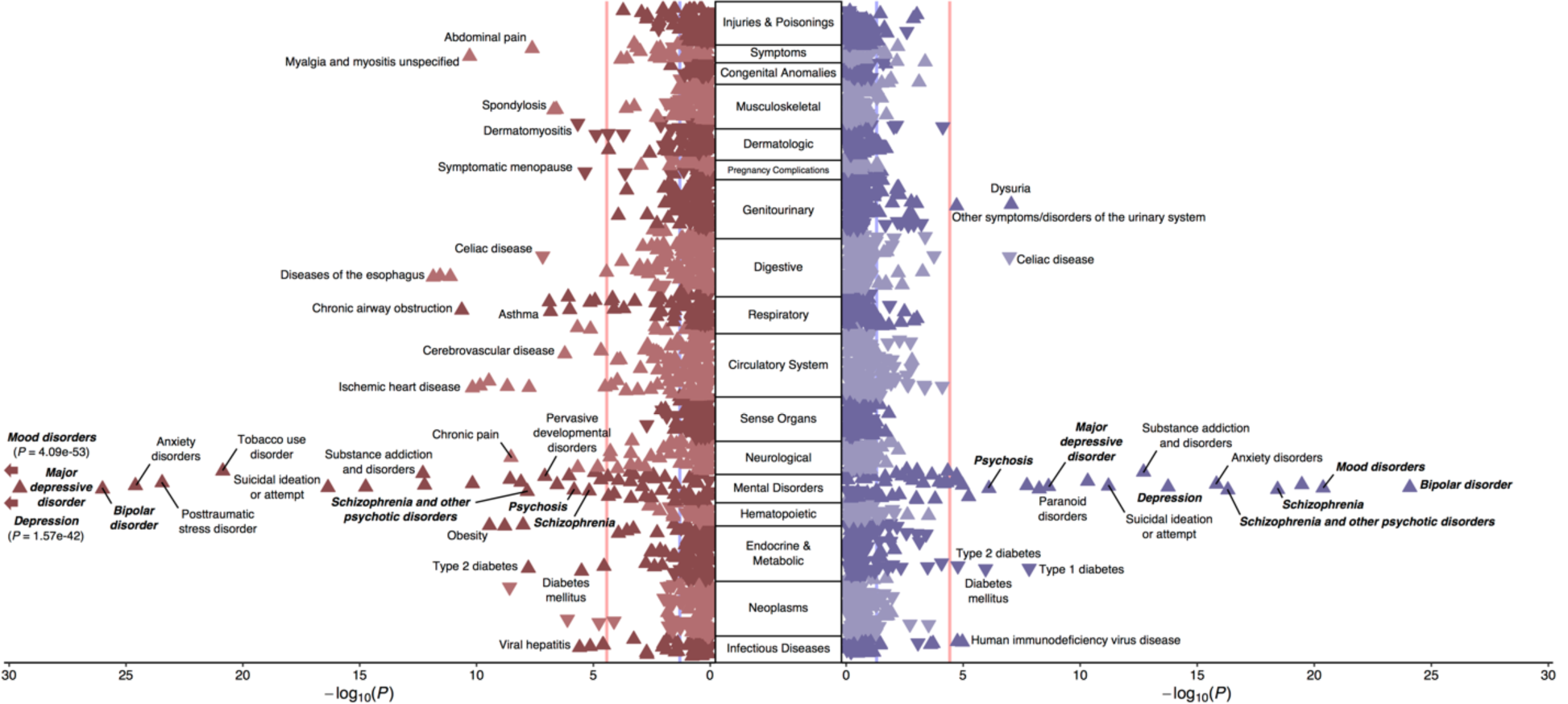
Phenome-wide association results for the two transdiagnostic latent genetic factors. Rotated Miami plot for (left) F1 and (right) F2, where the y-axis refers to the ICD-10 code category, the x-axis refers to the significance on a -log_10_ scale, the vertical light-red line denotes phenome-wide significance (*P* = 3.27e-5) following Bonferroni correction, and the vertical light-blue line marks nominal significance (*P* = .05). The direction of the triangle refers to the direction of effect. Phecodes closely resembling Genomic SEM model phenotypes are bolded and italicized for emphasis.

## Discussion

By jointly analyzing genome-wide data for eight psychiatric disorders and symptoms in a novel multivariate framework, we identified two distinct transdiagnostic factors that distinguished common forms of psychopathology related to mood disturbance versus rare forms of serious mental illness. Together, these factors explained approximately 80% of the genetic variance in mood and psychotic psychopathology, but were themselves only moderately correlated. Extensive biological annotation of these two transdiagnostic factors revealed clear differences between their factors in their underlying genetic architecture and biology. Further follow-up highlighted additional differences between the factors in their associations with human well-being and disease. Our results provide four critical insights into the genetic architecture of forms of psychopathology characterized by mood disturbance and psychosis.

First, we built on genomic investigations of the dimensional structure of certain forms of psychopathology, such as a large-scale study of the mood-disorder spectrum^23^, and identified two transdiagnostic factors that explain the vast majority of genetic variance in their constituent phenotypes. Perhaps surprisingly, variation in self-reported manic and psychotic symptoms is much more closely related to common forms of mood psychopathology (self-reported depressive symptoms, major depressive disorder, bipolar II disorder) than to psychiatric disorders characterized by severe mood disturbance and/or psychosis. Notably, we also find that the factor structure at the genetic level is different than the factor structure that we observe at the phenotypic level in the UK Biobank with similar indicators. This finding contrasts with what has been called the “phenotypic null hypothesis,” which states that genetic and phenotypic factor structures are expected to converge.^24^ Overall, these results illustrate how diagnostic boundaries, which are known to be problematic based on widespread phenotypic comorbidity, become even fuzzier at the genetic level of analysis.

Second, our multivariate association analyses identified 80 approximately independent loci associated with one of the transdiagnostic factors. Many of these genome-wide significant loci contain novel lead SNPs and map to genes that have not been previously associated with mood or psychotic psychopathology, such as *SIN3A*, which has been reported to be a key transcriptional regulator of cortical neurodevelopment, involved in neurogenesis and corticocortical projections in the developing mammalian brain^25^. Moreover, by employing multiple gene mapping techniques, we were also able to triangulate on novel genes associated with psychopathology, including *WDR73*, the causal gene in a rare recessive autosomal disorder characterized by severe encephalopathy, developmental delay, and neurocognitive impairment^26^. Associations such as these are particularly interesting in light of results suggesting that genes disrupted in Mendelian disorders are also dysregulated by non-coding variants in phenotypically-similar traits and disorders^27^. Furthermore, we build on the results of a large GWAS of eight psychiatric disorders^28^ by providing novel evidence of factor-specific pleiotropy (*i.e*., consistent effects across a factor’s constituent indicators) via *Q*_SNP_ results, which also identified several novel loci with significantly heterogeneous effects for bipolar I disorder, schizoaffective disorder, and schizophrenia.

Third, our extensive biological annotation revealed a marked divergence in the biology associated with the two transdiagnostic factors. While we find that the central nervous system is dually implicated at a broad systems-based level (*e.g*., non-specific enrichment of brain tissues), the biology associated with the two factors quickly diverges at more molecular levels of investigation. Via our novel approach to gene prioritization based on spatial transcriptomics, we identified two sets of factor-specific genes with contrasting developmental expression profiles. Specifically, we found that transcriptomically-prioritized genes associated with the factor broadly characterized by common mood disturbance (F1) exhibited *lower* expression levels during early prenatal periods, while transcriptomically-prioritized genes for the factor broadly characterized by rarer forms of serious mental illness (F2) exhibited *higher* expression levels during early prenatal periods. Notably, both of these trajectories identify the prenatal epoch as a critical developmental period related to psychopathology, albeit in different ways. These findings coalesce with and build upon previous studies that have begun to characterize developmental expression patterns of transdiagnostic genetic liabilities^29^. Here, we found that the two observed trajectories strongly resembled those of postnatal and prenatal inhibitory neuronal genes^21^, which have been implicated in the development of mood and psychotic disorders^30–32^.

Fourth, we found that the two factors differ substantially in their associations with human well-being and disease. Our results expand upon recent phenome-wide association studies of genetic risk for major depressive disorder^33^ and schizophrenia^34^, expanding the list of complex traits and medical phenotypes associated with mood and psychotic psychopathology. We also identified an interesting pattern of results in our genetic correlation and phenome-wide association analyses, where the factor comprising more common forms of mood disturbance (F1) had broader and often stronger negative associations with socioeconomic and health-related outcomes than the factor comprising rarer forms of serious mental illness (F2). This runs counter to associations often observed at the phenotypic level, where individuals diagnosed with more serious mental illnesses tend to face more severe impairments and consequences in these domains^35,36^.These results raise questions the potential ascertainment biases that affect genome-wide association studies. For example, clinically-ascertained samples of people with diagnosed psychiatric disorders (particularly when those disorders are rare and seriously impairing) are subject to different sources of selection, attrition, and non-response than population-based studies that utilize self-report surveys. Consider, for instance, that homeless and incarcerated individuals in Western countries are drastically more likely than the general population to meet diagnostic criteria for a serious mental illness^37,38^, but these socially-marginalized groups are less likely to have access to adequate mental health care or be included in medical research. This selective representation of psychopathology may induce collider bias and lead to misleading estimates of genetic association^39^. Indeed, cohort-level studies have already found that educational attainment polygenic scores are positively associated with research participation, while psychopathology polygenic scores are negatively associated^40,41^.

While we have taken many steps to address potential confounds, these major findings should be interpreted in light of several limitations. First, structural equation modeling does not reveal a “ground truth” about the nature of the phenotypes included in the analysis. Instead, it is a useful statistical framework for representing complex data structures, and latent factors are most appropriately considered as convenient statistical entities that explain the (co)variances of their indicators. As such, latent genetic factors are most useful as explanatory devices when accompanied by extensive biological annotation and follow-up as done in the present study. Second, the univariate GWASs are comprised of different samples with different measurement approaches and varying levels of power. However, we have made efforts to harmonize each of the GWASs used in Genomic SEM analyses (*e.g*., excluding self-rated measures from diagnostic phenotypes and vice-versa), and previous examination of these concerns suggest that the genetic factor structure is not biased by sample overlap or sample size differences^28,42^. Third, the univariate GWASs are comprised of different cohorts that may be subject to different sources of bias that cannot be fully quantified. Fourth, the current study focuses on forms of psychopathology that involve a wide variety of disturbances in mood and reality testing, but does not comprehensively sample the full range of psychiatric disorders. These results thus complement other transdiagnostic research studies that have illuminated how schizophrenia and bipolar I diverge genetically from other clinically-defined disorders, such as compulsive disorders and disorders of childhood^28^.

In conclusion, we have conducted a novel multivariate GWAS of multiple symptoms and disorders spanning mood and psychotic psychopathology. This analysis identified two transdiagnostic genetic liabilities operating quite distinctly from one another. Extensive biological annotation revealed contrasting genetic architectures that implicated prenatal neurodevelopment and neuronal function and regulation in markedly different ways. Given the degree of divergence between these two factors, future research is warranted to investigate the utility and appropriateness of even broader spectra of psychopathology (*e.g*., the *p* factor^43^) as explanatory devices at the level of molecular genetics. Collectively, our results suggest that the severity of mood and psychotic symptoms evident in severe psychiatric disorders might actually reflect a difference in kind, rather than merely in degree.

## Methods

### Genome-wide association analyses

#### Phenotype construction

M*plus*^44^ v8 was used to estimate person-specific thetas (*i.e*., factor scores) for three symptom domains: depression, mania, and psychosis. As each psychiatric phenotypes was assessed by four items, thetas were estimated via a multidimensional two-parameter probit model^45^, which allowed item-level responses across measurement occasions to be combined for correlated latent variables simultaneously. Furthermore, a combination of multiple imputation and Bayesian estimation with non-informative priors was used to maximally leverage all available responses for participants to minimize the impact of missing data. See Supplementary Section 1.1 for further description of the phenotypic modeling.

#### Univariate association analyses

BOLT-LMM^46^ v2.3.2 was used to conduct GWASs in the UK Biobank for three lifetime measures of psychiatric symptoms: depression, mania, and psychosis. This approach used a linear mixed model that included a genetic relationship matrix to estimate SNP effects, which offered improved control for population stratification and maximized power by accounting for relatedness among individuals. The first 40 principal components of ancestry computed with flashPCA2 (Supplementary Section 1.2), sex, birth year, sex-by-birth year interactions, and batch were included as covariates. EasyQC^47^ was used to perform extensive quality control on the GWAS summary statistics. The main objective of the quality control was to filter out rare and low-frequency SNPs, as well as SNPs that were not imputed well. Three main filters were imposed: (i) MAF < .005; (ii) imputation quality score < .9; (iii) unavailable in reference panel. Additional quality control procedures and filters are further described in Supplementary Section 1.3. The reference panel was a combination of the 1000 Genomes phase 3 v5 and UK10K, which has been described in a previous study^14^.

#### Summary statistics for psychiatric disorders

GWAS summary statistics for major depressive disorder, bipolar II disorder, bipolar I disorder, schizoaffective disorder, and schizophrenia were obtained from the Psychiatric Genomics Consortium. Quality control was performed in the original studies, but additional filters were applied as necessary to harmonize with the quality control pipeline in the present study. All GWAS samples were restricted to European ancestry to minimize the potential influence of population stratification.

#### Multivariate association analyses

Genomic SEM^42^ v0.0.2 was used to conduct multivariate GWAS based on eight phenotypes: depressive symptoms, manic symptoms, psychotic symptoms, major depressive disorder, bipolar II disorder, bipolar I disorder, schizoaffective disorder, and schizophrenia (see Table 1 for overview). Following identification of the confirmatory factor model that best explained the observed genetic covariances among the phenotypes, Genomic SEM was used to estimate the individual SNP effects on each latent factor in the model. Note that Genomic SEM is unbiased in the presence of varying and unknown sample overlap across the contributing GWAS samples, as the cross-trait intercepts estimated via multivariable LD score regression are used to estimate (and account for) sample overlap and phenotypic correlation.

Effective sample size (*N*_*eff*_) for each latent factor was estimated as 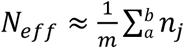, where, is the number of SNPs in the GWAS, + is the lower MAF threshold for inclusion in the calculation (here, 10%), * the upper limit (here, 40%), and *n*_*j*_ is the effective sample size for SNP *j*, which is calculated as 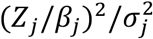. Q_SNP_ tests were used to evaluate whether SNP effects on the latent factors were driven by heterogenous effects across constituent phenotypes. Further description of multivariate association analyses and Q_SNP_ tests is provided in Supplementary Sections 2.4 and 2.5, respectively.

### Genomic structural equation modeling

#### Genetic correlations among study phenotypes

LD score regression^48^ v1.0.1 was used to estimate genetic correlations between all pairwise combinations of the eight study phenotypes. Standard procedures and best practices for LD score regression were followed (*e.g*., restricting to HapMap3^49^ SNPs with a minor allele frequency ≥ .01). Default parameters were used for the three new GWASs of psychiatric symptoms. For the existing GWASs of psychiatric disorders, parameters (*e.g*., sample prevalence, population prevalence) were defined as outlined in the original studies. A hierarchical clustering algorithm was applied to the final genetic correlation matrix to guide factor selection in the exploratory factor analysis. Although the original LD score regression software was used for this preliminary analysis, the multivariable version of LD score regression employed by Genomic SEM was used for all subsequent analyses. Please note that these software produce estimates that are effectively identical.

#### Exploratory factor analysis

The stats R package was used to conduct an EFA of the genetic correlations among the eight study phenotypes. Specifically, the *factanal* function was used to conduct an EFA with promax rotation on the standardized *S* matrix derived from the multivariable version of LD score regression employed by Genomic SEM. This enabled an empirical assessment of (i) the number of latent factors that best explained the multivariate genetic architecture observed among the set of study phenotypes (*i.e*., the number of transdiagnostic liabilities present), and (ii) how constituent phenotypes load onto separable latent factors. As suggested by the hierarchical clustering algorithm, two factors were extracted that optimally accounted for shared variation among sets of the observed variables. Results from this analysis were subsequently used to guide construction of the confirmatory factor models. A brief overview of factor analysis is provided in Supplementary Section 2.2.

#### Confirmatory factor analysis

Genomic SEM was used to test whether a common factor model or a correlated factors model best fit the data via CFA, where fit reflects the degree to which the specified latent variable structure adequately explains the observed covariances among the set of observed variables. Parameter estimates were derived using weighted least squares estimation. Model fit was assessed using conventional indices in structural equation modeling: the model χ^2^ statistic, the Akaike information criterion (AIC), the comparative fit index (CFI), and the standardized root mean square residual (SRMR). All fit indices retain their standard interpretations within a Genomic SEM framework. However the model χ^2^ statistic is best used as a comparative measure of fit to evaluate competing models rather than a measure of statistical significance given the sensitivity of model χ^2^ to sample size, which is comparatively extremely large for GWAS samples. For CFI and SRMR, values greater than .90 and less than .08, respectively, were considered reflective of good model fit^50^. Further description of structural equation modeling and confirmatory factor analysis are provided in Supplementary Section 2.3.

### Heritability analyses

#### Heritability for observed and latent phenotypes

LD score regression was used to estimate heritabilities for the three novel univariate GWASs, as well as the two novel multivariate GWASs. Standard procedures and best practices for LD score regression were followed. As there is no phenotypic variance for latent genetic factors modeled in Genomic SEM, heritability is more accurately referred to as genetic variance for F1 and F2. Furthermore, as genetic variance estimates are influenced by the heritabilities of constituent phenotypes and the metric of the latent genetic factor, estimates for F1 and F2 should only be interpreted in the context of the present study.

#### Local heritability and genetic correlations

HESS^51^ and its bivariate extension, *ρ*-HESS^52^, were used to estimate local genetic variance, local genetic covariance, and the proportion of the genome that contributes to the total genetic covariance for F1 and F2. For each factor, HESS was first used to estimate local genetic variance and covariance across 1,698 approximately LD-independent contiguous genomic partitions, averaging 1.5 Mb per partition. The European samples from the 1000 Genomes Project Phase 3v5^53^ (*n* = 503) were used as a reference panel for these analyses. Independent genomic partitions were then ranked by their absolute genetic covariance, and the percentage that accounted for 80% of the total genetic covariance between F1 and F2 was used to further quantify genetic overlap between F1 and F2^54^.

### Gene mapping and identification

The FUMA^55^ SNP2GENE pipeline was used to apply a standard clumping algorithm that identified associated genomic loci, lead SNPs within loci, and all independent significant SNPs within loci. The European samples from the 1000 Genomes Project Phase 3v5 (*n* = 503) were used as a reference panel for LD. FUMA was also used to employ an ensemble of methods to identify putative risk genes for the univariate and multivariate GWAS phenotypes. Specifically, FUMA v1.3.5e was used to conduct positional, eQTL, and chromatin interaction mapping to identify risk-conferring genes that map to genome-wide significant loci. Default parameters were used for each of these analyses. ANNOVAR annotations^56^ were used for positional mapping, the Genotype-Tissue Expression (GTEx) v8 brain dataset^57^ was used as the reference tissue data for eQTL mapping, and Hi-C data from adult and fetal human brain samples^58^ was used to examine enhancer-promoter and promoter-promoter chromatin interactions. The FUMA GENE2FUNC pipeline was used to identify overlap between identified genes and biological gene sets as catalogued by MolSigDB v7.0, as well as previous hits in GWAS Catalog (https://www.ebi.ac.uk/gwas/).

Two additional methods were employed to identify putative risk genes based on genome-wide summary statistics: and MAGMA^59^ and S-PrediXcan^60^. The former was used to identify functionally expressed genes via joint analysis of SNP effects and eQTL expression effects, and the latter was used to calculate gene-based association statistics. Both methods are described in the following section.

### Gene-based association and enrichment analyses

MAGMA v1.07, a bioinformatics software for gene-based biological annotation, was used to conduct gene association, gene set enrichment, and gene property analyses for all novel study phenotypes. Default MAGMA parameters were employed and standard procedures were followed for gene-based association analyses based on summary statistics. For gene-level association analyses, test statistics were computed using a window of 10kb around the gene of interest for all novel GWAS phenotypes. MAGMA was then used to conduct competitive gene-set enrichment and gene property analyses based on the gene-level *P* values produced in the association analyses. These analyses tested whether genes within an annotated set are more strongly associated with the phenotype of interest than other genes. For gene set enrichment analyses, up to 15,481 gene sets catalogued in MolSigDB v7.0 were tested, which corresponded to 7,341 biological processes, 1,000 cellular components, 1,642 molecular functions, and 5,496 expertly curated gene sets broadly related to biological pathways and processes. For the gene property analyses, 54 tissues from the GTEx v8 dataset were tested. Bonferroni-corrected thresholds of *P* ≤ 3.23e-6 and *P* ≤ 9.26e-4 were used to determine significance for gene sets and tissues, respectively.

S-PrediXcan v0.6.2 was used (i) to predict gene expression levels in brain tissues, and (ii) to test whether predicted gene expression correlated with either transdiagnostic factor. Tissue weights were computed using reference data from the GTEx v8 dataset. GWAS summary statistics for F1 and F2, the reference transcriptomic data, and covariance matrices for the SNPs within each gene model were included as input data. Thirteen brain tissues were tested: anterior cingulate cortex, amygdala, caudate basal ganglia, cerebellar hemisphere, cerebellum, cortex, frontal cortex, hippocampus, hypothalamus, nucleus accumbens basal ganglia, putamen basal ganglia, spinal cord and substantia nigra. A Bonferonni-corrected threshold of *P* ≤ 8.97e-7 was established for transcriptome-wide significance, which corrected for 55,753 gene-based tests.

### Genetic correlation analyses

Genomic SEM was used to estimate genetic correlations and partial genetic correlations between latent factors of psychopathology and other phenotypes of interest. Specifically, genetic correlations were estimated for two broad sets of phenotypes: (i) morphological features of the human brain, and (ii) complex traits related to human health and well-being. Summary statistics for 101 neuroimaging phenotypes^61^ (cortical and subcortical gray matter volumes, ventricular volumes, and global measures of brain volume) were downloaded from https://github.com/BIG-S2/GWAS. Summary statistics for 92 phenotypes broadly related to various domains of human health and well-being were downloaded from various online sources, using download links from GWAS Atlas^62^ whenever possible. All summary statistics were cleaned and processed using the *munge* function of Genomic SEM, retaining all HapMap3 SNPs outside of the major histocompatibility complex regions with an allele frequency ≥ .01. A Bonferroni correction was applied within each family of tests to adjust *P* values for multiple comparisons (*P* ≤ 4.95e-4 for neuroimaging phenotypes; *P* ≤ 5.43e-4 for complex traits).

### Spatiotemporal transcriptomic analyses

Microarray gene expression data from the Allen Human Brain Atlas (AHBA)^63^ were downloaded from https://human.brain-map.org/static/download, and subsequently aligned to the Desikan-Killiany-Tourville atlas (*N* = 62 cortical brain regions)^64^ for spatial compatibility with the cortical neuroimaging phenotypes^65^. Spatial correlation coefficients (Spearman’s *ρ*) were computed for each of 20,647 genes compared against the -log_10_ *P* values from F1 and F2. To examine the developmental trajectories of the F1 and F2 gene sets (positive *Z*-scores of AHBA correlation coefficients, *P* < .05), weighted gene correlation network analysis^66^ was used to estimate eigengene values (*i.e*., gene set expression) for these gene sets in the Brainspan dataset, treating each factor-specific gene set as a module. These expression values were then plotted as function of time, using a non-parametric LOESS curve line-of-best-fit to characterize developmental expression trajectories for F1 and F2, which indicated that the prioritized gene sets for each transdiagnostic factor are differentially expressed in pre- and postnatal cortical tissue. Evaluation of cell-type-specific gene sets was performed as above, using available data from a recent cell-specific sequencing study in adult human brain tissue^21^.

### Phenome-wide polygenic prediction

PRS-CS^67^ and PLINK^68^ v1.9 were used to calculate polygenic scores for the transdiagnostic latent genetic factors, F1 and F2. PRS-CS, a Bayesian polygenic prediction method, was used to apply a continuous shrinkage prior to SNP effect estimates and infer posterior SNP weights using GWAS summary statistics for F1 and F2 and an external reference panel to model LD. In the present study, PRS-CS was used to adjust weights for 1,027,871 SNPs typed on both the 1000 Genomes Project Phase 3v5 and the HapMap3 reference panels with a minor allele frequency ≥ .01. The European samples from the 1000 Genomes Project Phase 3v5 (*n* = 503) were used as a reference panel for LD. PLINK was then used to calculate polygenic scores for each individual by summing all included variants weighted by the inferred posterior effect size for the effect allele, and converting that value to a Z-score for each participant within the prediction sample.

The genotyped BioVU sample (*N* = 66,915) was used to test for associations between polygenic scores for F1 and F2 and a wide array of medical phenotypes. Genotyping and quality control for this sample have been described elsewhere. Case-control medical phenotypes, also referred to as “phecodes,” were constructed from International Classification of Disease (ICD) diagnostic codes in participant electronic health record data. Two instances of an ICD diagnostic code were required to be present to be classified as a case for a given phecode, and 50 cases were required for a phecode to be analyzed. A total of 1,335 phecodes were included in the phenome-wide association analyses. The PheWAS R package was used to conduct phenome-wide association analyses. A logistic regression model was fit to each of 1,335 case/control phenotypes to estimate the odds of each diagnosis given the polygenic scores for F1 and F2. Sex, median age of the longitudinal electronic health record measurements, and the top 10 principal components of ancestry were included as covariates. A Bonferroni-corrected threshold of at *P* ≤ 3.74e-5 was established for phenome-wide significance.

## Acknowledgments

K.P.H. and E.M.T.D. are supported by Jacobs Foundation Research Fellowships, and are Faculty Research Associates of the Population Research Center at the University of Texas at Austin, which is supported by a grant 5-R24-HD042849 from the Eunice Kennedy Shriver National Institute of Child Health and Human Development (NICHD). Research by K.P.H. and E.M.T.D. is further supported by NICHD grant R01-HD083613. P.D.K was supported by an ERC consolidator grant (647648 EdGe). This research was conducted using the UK Biobank Resource under Application Number 11425, and with the support and collaboration from all investigators who make up the Bipolar Disorder Working Group of the PGC (full list in the Supplementary Note). We would like to thank the many studies that made these consortia possible, the researchers involved, and the participants in those studies, without whom this effort would not be possible. We would also like to thank the research participants and employees of 23andMe for making this work possible.

## Author contributions

T.T.M., P.D.K., and K.P.H. conceived and designed the study. P.D.K. and K.P.H. oversaw the study. T.T.M. and K.P.H. led the writing of the manuscript, with substantive contributions from P.D.K. and M.C.K. A.A.P., E.M.T-D., and K.S.K. provided valuable feedback on the framing and interpretation of the results. T.T.M. was the lead analyst, responsible for conducting genome-wide association studies, quality control, genetic correlations, multivariate analyses with Genomic SEM, and biological annotation, with assistance from R.K.L., A.D.G., S.S-R., and J.S. T.T.M. prepared the data for analysis, with assistance from R.K.L., A.O., R.D.V., and S.F.W.M. S.S-R. performed the phenome-wide association study with assistance from L.K.D. T.T.M. prepared the figures and tables. Investigators from the Bipolar Disorder Working Group of the PGC contributed data for bipolar I, bipolar II, and schizoaffective disorder. All authors provided valuable feedback and advice during preparation of the manuscript.

## Data availability

All genetic and phenotypic data from UK Biobank are available via their standard data access procedure, described at https://www.ukbiobank.ac.uk/register-apply. Summary statistics for F1 and F2 are available at [link to data repository to be provided upon publication]. Similarly, summary statistics for symptoms of depression, mania, and psychosis are available at [link to data repository to be provided upon publication]. Summary statistics for major depressive disorder, bipolar I, bipolar II, schizoaffective disorder, and schizophrenia can be download or requested at https://www.med.unc.edu/pgc/download-results/. 23andMe summary statistics are made available through 23andMe to qualified researchers under an agreement with 23andMe that protects the privacy of 23andMe participants. Please visit https://research.23andme.com/collaborate/ for more information.

