## Supplementary Note for "Multivariate GWAS of psychiatric disorders and their cardinal symptoms reveal two dimensions of cross-cutting genetic liabilities"

#### Affiliations:

Eli A Stahl1,2,3†, Gerome Breen4,5†, Andreas J Forstner6,7,8,9,10†, Andrew McQuillin11†, Stephan Ripke12,13,14†, Vassily Trubetskoy13, Manuel Mattheisen15,16,17,18,19, Yunpeng Wang20,21, Jonathan R I Coleman4,5, Héléna A Gaspar4,5, Christiaan A de Leeuw22, Stacy Steinberg23, Jennifer M Whitehead Pavlides24, Maciej Trzaskowski25, Enda M Byrne25, Tune H Pers3,26, Peter A Holmans27, Alexander L Richards27, Liam Abbott12, Esben Agerbo19,28,29, Huda Akil30, Diego Albani31, Ney Alliey-Rodriguez32, Thomas D Als15,16,19, Adebayo Anjorin33, Verneri Antilla14, Swapnil Awasthi13, Judith A Badner34, Marie Bækvad-Hansen19,35, Jack D Barchas36, Nicholas Bass11, Michael Bauer37, Richard Belliveau12, Sarah E Bergen38, Carsten Bøcker Pedersen19,28,29, Erlend Bøen39, Marco P. Boks40, James Boocock41, Monika Budde42, William Bunney43, Margit Burmeister44, Jonas Bybjerg-Grauholm19,35, William Byerley45, Miquel Casas46,47,48,49, Felecia Cerrato12, Pablo Cervantes50, Kimberly Chambert12, Alexander W Charney2, Danfeng Chen12, Claire Churchhouse12,14, Toni-Kim Clarke51, William Coryell52, David W Craig53, Cristiana Cruceanu50,54, David Curtis55,56, Piotr M Czerski57, Anders M Dale58,59,60,61, Simone de Jong4,5, Franziska Degenhardt8, Jurgen Del-Favero62, J Raymond DePaulo63, Srdjan Djurovic64,65, Amanda L Dobbyn1,2, Ashley Dumont12, Torbjørn Elvsåshagen66,67, Valentina Escott-Price27, Chun Chieh Fan61, Sascha B Fischer6,10, Matthew Flickinger68, Tatiana M Foroud69, Liz Forty27, Josef Frank70, Christine Fraser27, Nelson B Freimer71, Louise Frisén72,73,74, Katrin Gade42,75, Diane Gage12, Julie Garnham76, Claudia Giambartolomei206, Marianne Giørtz Pedersen19,28,29, Jaqueline Goldstein12, Scott D Gordon77, Katherine Gordon-Smith78, Elaine K Green79, Melissa J Green80,133, Tiffany A Greenwood60, Jakob Grove15,16,19,81, Weihua Guan82, José Guzman-Parra83, Marian L Hamshere27, Martin Hautzinger84, Urs Heilbronner42, Stefan Herms6,8,10, Maria Hipolito85, Per Hoffmann6,8,10, Dominic Holland58,86, Laura Huckins1,2, Stéphane Jamain87,88, Jessica S Johnson1,2, Radhika Kandaswamy4, Robert Karlsson38, James L Kennedy89,90,91,92, Sarah Kittel-Schneider93, James A Knowles94,95, Manolis Kogevinas96, Anna C Koller8, Ralph Kupka97,98,99, Catharina Lavebratt72, Jacob Lawrence100, William B Lawson85, Markus Leber101, Phil H Lee12,14,102, Shawn E Levy103, Jun Z Li104, Chunyu Liu105, Susanne Lucae106, Anna Maaser8, Donald J MacIntyre107,108, Pamela B Mahon63,109, Wolfgang Maier110, Lina Martinsson73, Steve McCarroll12,111, Peter McGuffin4, Melvin G McInnis112, James D McKay113, Helena Medeiros95, Sarah E Medland77, Fan Meng30,112, Lili Milani114, Grant W Montgomery25, Derek W Morris115,116, Thomas W Mühleisen6,117, Niamh Mullins4, Hoang Nguyen1,2, Caroline M Nievergelt60,118, Annelie Nordin Adolfsson119, Evaristus A Nwulia85, Claire O'Donovan76, Loes M Olde Loohuis71, Anil P S Ori71, Lilijana Oruc120, Urban Ösby121, Roy H Perlis122,123, Amy Perry78, Andrea Pfennig37, James B Potash63, Shaun M Purcell2,109, Eline J Regeer124, Andreas Reif93, Céline S Reinbold6,10, John P Rice125, Fabio Rivas83, Margarita Rivera4,126, Panos Roussos1,2,127, Douglas M Ruderfer128, Euijung Ryu129, Cristina Sánchez-Mora46,47,49, Alan F Schatzberg130, William A Scheftner131, Nicholas J Schork132, Cynthia Shannon Weickert80,133, Tatyana Shehktman60, Paul D Shilling60, Engilbert Sigurdsson134, Claire Slaney76, Olav B Smeland135,136, Janet L Sobell137, Christine Søholm Hansen19,35, Anne T Spijker138, David St Clair139, Michael Steffens140, John S Strauss91,141, Fabian Streit70, Jana Strohmaier70, Szabolcs Szelinger142, Robert C Thompson112, Thorgeir E Thorgeirsson23, Jens Treutlein70, Helmut Vedder143, Weiqing Wang1,2, Stanley J Watson112, Thomas W Weickert80,133, Stephanie H Witt70, Simon Xi144, Wei Xu145,146, Allan H Young147, Peter Zandi148, Peng Zhang149, Sebastian Zöllner112, eQTLGen Consortium, BIOS Consortium, Rolf Adolfsson119, Ingrid Agartz17,39,150, Martin Alda76,151, Lena Backlund73, Bernhard T Baune152,158, Frank Bellivier153,154,155,156, Wade H Berrettini157, Joanna M Biernacka129, Douglas H R Blackwood51, Michael Boehnke68, Anders D Børglum15,16,19, Aiden Corvin116, Nicholas Craddock27, Mark J Daly12,14, Udo Dannlowski158, Tõnu Esko3,111,114,159, Bruno Etain153,155,156,160, Mark Frye161, Janice M Fullerton133,162, Elliot S Gershon32,163, Michael Gill116, Fernando Goes63, Maria Grigoroiu-Serbanescu164, Joanna Hauser57, David M Hougaard19,35, Christina M Hultman38, Ian Jones27, Lisa A Jones78, René S Kahn2,40, George Kirov27, Mikael Landén38,165, Marion Leboyer88,153,166, Cathryn M Lewis4,5,167, Qingqin S Li168, Jolanta Lissowska169, Nicholas G Martin77,170, Fermin Mayoral83, Susan L McElroy171, Andrew M McIntosh51,172, Francis J McMahon173, Ingrid Melle174,175, Andres Metspalu114,176, Philip B Mitchell80, Gunnar Morken177,178, Ole Mors19,179, Preben Bo Mortensen15,19,28,29, Bertram Müller-Myhsok54,180,181, Richard M Myers103, Benjamin M Neale3,12,14, Vishwajit Nimgaonkar182, Merete Nordentoft19,183, Markus M Nöthen8, Michael C O'Donovan27, Ketil J Oedegaard184,185, Michael J Owen27, Sara A Paciga186, Carlos Pato95,187, Michele T Pato95, Danielle Posthuma22,188, Josep Antoni Ramos-Quiroga46,47,48,49, Marta Ribasés46,47,49, Marcella Rietschel70, Guy A Rouleau189,190, Martin Schalling72, Peter R Schofield133,162, Thomas G Schulze42,63,70,75,173, Alessandro Serretti191, Jordan W Smoller12,192,193, Hreinn Stefansson23, Kari Stefansson23,194, Eystein Stordal195,196, Patrick F Sullivan38,197,198, Gustavo Turecki199, Arne E Vaaler200, Eduard Vieta201, John B Vincent141, Thomas Werge19,202,203, John I Nurnberger204, Naomi R Wray24,25, Arianna Di Florio27,198, Howard J Edenberg205, Sven Cichon6,8,10,117, Roel A Ophoff40,41,71, Laura J Scott68, Ole A Andreassen135,136, John Kelsoe60*, Pamela Sklar1,2*^

† Equal contribution
* Co-senior investigators
^ Deceased

Affiliations:

1 Department of Genetics and Genomic Sciences, Icahn School of Medicine at Mount Sinai, New York, NY, US
2 Department of Psychiatry, Icahn School of Medicine at Mount Sinai, New York, NY, US
3 Medical and Population Genetics, Broad Institute, Cambridge, MA, US
4 MRC Social, Genetic and Developmental Psychiatry Centre, King's College London, London, GB
5 NIHR BRC for Mental Health, King's College London, London, GB
6 Department of Biomedicine, University of Basel, Basel, CH
7 Department of Psychiatry (UPK), University of Basel, Basel, CH
8 Institute of Human Genetics, University of Bonn, School of Medicine & University Hospital Bonn, Bonn, DE
9 Centre for Human Genetics, University of Marburg, Marburg, DE
10 Institute of Medical Genetics and Pathology, University Hospital Basel, Basel, CH
11 Division of Psychiatry, University College London, London, GB
12 Stanley Center for Psychiatric Research, Broad Institute, Cambridge, MA, US
13 Department of Psychiatry and Psychotherapy, Charité - Universitätsmedizin, Berlin, DE
14 Analytic and Translational Genetics Unit, Massachusetts General Hospital, Boston, MA, US
15 iSEQ, Center for Integrative Sequencing, Aarhus University, Aarhus, DK
16 Department of Biomedicine - Human Genetics, Aarhus University, Aarhus, DK
17 Department of Clinical Neuroscience, Centre for Psychiatry Research, Karolinska Institutet, Stockholm, SE
18 Department of Psychiatry, Psychosomatics and Psychotherapy, Center of Mental Health, University Hospital Würzburg, Würzburg, DE
19 iPSYCH, The Lundbeck Foundation Initiative for Integrative Psychiatric Research, DK
20 Institute of Biological Psychiatry, Mental Health Centre Sct. Hans, Copenhagen, DK
21 Institute of Clinical Medicine, University of Oslo, Oslo, NO
22 Department of Complex Trait Genetics, Center for Neurogenomics and Cognitive Research, Amsterdam Neuroscience, Vrije Universiteit Amsterdam, Amsterdam, NL
23 deCODE Genetics / Amgen, Reykjavik, IS
24 Queensland Brain Institute, The University of Queensland, Brisbane, QLD, AU
25 Institute for Molecular Bioscience, The University of Queensland, Brisbane, QLD, AU
26 Division of Endocrinology and Center for Basic and Translational Obesity Research, Boston Children's Hospital, Boston, MA, US
27 Medical Research Council Centre for Neuropsychiatric Genetics and Genomics, Division of Psychological Medicine and Clinical Neurosciences, Cardiff University, Cardiff, GB
28 National Centre for Register-Based Research, Aarhus University, Aarhus, DK
29 Centre for Integrated Register-based Research, Aarhus University, Aarhus, DK
30 Molecular & Behavioral Neuroscience Institute, University of Michigan, Ann Arbor, MI, US
31 Department of Neuroscience, IRCCS - Istituto Di Ricerche Farmacologiche Mario Negri, Milan, IT
32 Department of Psychiatry and Behavioral Neuroscience, University of Chicago, Chicago, IL, US
33 Psychiatry, Berkshire Healthcare NHS Foundation Trust, Bracknell, GB
34 Psychiatry, Rush University Medical Center, Chicago, IL, US
35 Center for Neonatal Screening, Department for Congenital Disorders, Statens Serum Institut, Copenhagen, DK
36 Department of Psychiatry, Weill Cornell Medical College, New York, NY, US
37 Department of Psychiatry and Psychotherapy, University Hospital Carl Gustav Carus, Technische Universität Dresden, Dresden, DE
38 Department of Medical Epidemiology and Biostatistics, Karolinska Institutet, Stockholm, SE
39 Department of Psychiatric Research, Diakonhjemmet Hospital, Oslo, NO
40 Psychiatry, UMC Utrecht Brain Center Rudolf Magnus, Utrecht, NL
41 Human Genetics, University of California Los Angeles, Los Angeles, CA, US
42 Institute of Psychiatric Phenomics and Genomics (IPPG), University Hospital, LMU Munich, Munich, DE
43 Department of Psychiatry and Human Behavior, University of California, Irvine, Irvine, CA, US
44 Molecular & Behavioral Neuroscience Institute and Department of Computational Medicine & Bioinformatics, University of Michigan, Ann Arbor, MI, US
45 Psychiatry, University of California San Francisco, San Francisco, CA, US
46 Instituto de Salud Carlos III, Biomedical Network Research Centre on Mental Health (CIBERSAM), Madrid, ES
47 Department of Psychiatry, Hospital Universitari Vall d´Hebron, Barcelona, ES
48 Department of Psychiatry and Forensic Medicine, Universitat Autònoma de Barcelona, Barcelona, ES
49 Psychiatric Genetics Unit, Group of Psychiatry Mental Health and Addictions, Vall d´Hebron Research Institut (VHIR), Universitat Autònoma de Barcelona, Barcelona, ES
50 Department of Psychiatry, Mood Disorders Program, McGill University Health Center, Montreal, QC, CA
51 Division of Psychiatry, University of Edinburgh, Edinburgh, GB

52 University of Iowa Hospitals and Clinics, Iowa City, IA, US
53 Translational Genomics, USC, Phoenix, AZ, US
54 Department of Translational Research in Psychiatry, Max Planck Institute of Psychiatry, Munich, DE
55 Centre for Psychiatry, Queen Mary University of London, London, GB
56 UCL Genetics Institute, University College London, London, GB
57 Department of Psychiatry, Laboratory of Psychiatric Genetics, Poznan University of Medical Sciences, Poznan, PL
58 Department of Neurosciences, University of California San Diego, La Jolla, CA, US
59 Department of Radiology, University of California San Diego, La Jolla, CA, US
60 Department of Psychiatry, University of California San Diego, La Jolla, CA, US
61 Department of Cognitive Science, University of California San Diego, La Jolla, CA, US
62 Applied Molecular Genomics Unit, VIB Department of Molecular Genetics, University of Antwerp, Antwerp, Belgium
63 Department of Psychiatry and Behavioral Sciences, Johns Hopkins University School of Medicine, Baltimore, MD, US
64 Department of Medical Genetics, Oslo University Hospital Ullevål, Oslo, NO
65 NORMENT, KG Jebsen Centre for Psychosis Research, Department of Clinical Science, University of Bergen, Bergen, NO
66 Department of Neurology, Oslo University Hospital, Oslo, NO
67 NORMENT, KG Jebsen Centre for Psychosis Research, Oslo University Hospital, Oslo, NO
68 Center for Statistical Genetics and Department of Biostatistics, University of Michigan, Ann Arbor, MI, US
69 Department of Medical & Molecular Genetics, Indiana University, Indianapolis, IN, US
70 Department of Genetic Epidemiology in Psychiatry, Central Institute of Mental Health, Medical Faculty Mannheim, Heidelberg University, Mannheim, DE
71 Center for Neurobehavioral Genetics, University of California Los Angeles, Los Angeles, CA, US
72 Department of Molecular Medicine and Surgery, Karolinska Institutet and Center for Molecular Medicine, Karolinska University Hospital, Stockholm, SE
73 Department of Clinical Neuroscience, Karolinska Institutet and Center for Molecular Medicine, Karolinska University Hospital, Stockholm, SE
74 Child and Adolescent Psychiatry Research Center, Stockholm, SE
75 Department of Psychiatry and Psychotherapy, University Medical Center Göttingen, Göttingen, DE
76 Department of Psychiatry, Dalhousie University, Halifax, NS, CA
77 Genetics and Computational Biology, QIMR Berghofer Medical Research Institute, Brisbane, QLD, AU
78 Department of Psychological Medicine, University of Worcester, Worcester, GB
79 School of Biomedical Sciences, Plymouth University Peninsula Schools of Medicine and Dentistry, University of Plymouth, Plymouth, GB
80 School of Psychiatry, University of New South Wales, Sydney, NSW, AU
81 Bioinformatics Research Centre, Aarhus University, Aarhus, DK
82 Biostatistics, University of Minnesota System, Minneapolis, MN, US
83 Mental Health Department, University Regional Hospital, Biomedicine Institute (IBIMA), Málaga, ES
84 Department of Psychology, Eberhard Karls Universität Tübingen, Tubingen, DE
85 Department of Psychiatry and Behavioral Sciences, Howard University Hospital, Washington, DC, US
86 Center for Multimodal Imaging and Genetics, University of California San Diego, La Jolla, CA, US
87 Psychiatrie Translationnelle, Inserm U955, Créteil, FR
88 Faculté de Médecine, Université Paris Est, Créteil, FR
89 Campbell Family Mental Health Research Institute, Centre for Addiction and Mental Health, Toronto, ON, CA
90 Neurogenetics Section, Centre for Addiction and Mental Health, Toronto, ON, CA
91 Department of Psychiatry, University of Toronto, Toronto, ON, CA
92 Institute of Medical Sciences, University of Toronto, Toronto, ON, CA
93 Department of Psychiatry, Psychosomatic Medicine and Psychotherapy, University Hospital Frankfurt, Frankfurt am Main, DE
94 Cell Biology, SUNY Downstate Medical Center College of Medicine, Brooklyn, NY, US
95 Institute for Genomic Health, SUNY Downstate Medical Center College of Medicine, Brooklyn, NY, US
96 ISGlobal, Barcelona, ES
97 Psychiatry, Altrecht, Utrecht, NL
98 Psychiatry, GGZ inGeest, Amsterdam, NL
99 Psychiatry, VU medisch centrum, Amsterdam, NL
100 Psychiatry, North East London NHS Foundation Trust, Ilford, GB
101 Clinic for Psychiatry and Psychotherapy, University Hospital Cologne, Cologne, DE
102 Psychiatric and Neurodevelopmental Genetics Unit, Massachusetts General Hospital, Boston, MA, US
103 HudsonAlpha Institute for Biotechnology, Huntsville, AL, US
104 Department of Human Genetics, University of Michigan, Ann Arbor, MI, US
105 Psychiatry, University of Illinois at Chicago College of Medicine, Chicago, IL, US
106 Max Planck Institute of Psychiatry, Munich, DE
107 Mental Health, NHS 24, Glasgow, GB

108 Division of Psychiatry, Centre for Clinical Brain Sciences, University of Edinburgh, Edinburgh, GB
109 Psychiatry, Brigham and Women's Hospital, Boston, MA, US
110 Department of Psychiatry and Psychotherapy, University of Bonn, Bonn, DE
111 Department of Genetics, Harvard Medical School, Boston, MA, US
112 Department of Psychiatry, University of Michigan, Ann Arbor, MI, US
113 Genetic Cancer Susceptibility Group, International Agency for Research on Cancer, Lyon, FR
114 Estonian Genome Center, University of Tartu, Tartu, EE
115 Discipline of Biochemistry, Neuroimaging and Cognitive Genomics (NICOG) Centre, National University of Ireland, Galway, Galway, IE
116 Neuropsychiatric Genetics Research Group, Dept of Psychiatry and Trinity Translational Medicine Institute, Trinity College Dublin, Dublin, IE
117 Institute of Neuroscience and Medicine (INM-1), Research Centre Jülich, Jülich, DE
118 Research/Psychiatry, Veterans Affairs San Diego Healthcare System, San Diego, CA, US
119 Department of Clinical Sciences, Psychiatry, Umeå University Medical Faculty, Umeå, SE
120 Department of Clinical Psychiatry, Psychiatry Clinic, Clinical Center University of Sarajevo, Sarajevo, BA
121 Department of Neurobiology, Care sciences, and Society, Karolinska Institutet and Center for Molecular Medicine, Karolinska University Hospital, Stockholm, SE
122 Psychiatry, Harvard Medical School, Boston, MA, US
123 Division of Clinical Research, Massachusetts General Hospital, Boston, MA, US
124 Outpatient Clinic for Bipolar Disorder, Altrecht, Utrecht, NL
125 Department of Psychiatry, Washington University in Saint Louis, Saint Louis, MO, US
126 Department of Biochemistry and Molecular Biology II, Institute of Neurosciences, Center for Biomedical Research, University of Granada, Granada, ES
127 Department of Neuroscience, Icahn School of Medicine at Mount Sinai, New York, NY, US
128 Medicine, Psychiatry, Biomedical Informatics, Vanderbilt University Medical Center, Nashville, TN, US
129 Department of Health Sciences Research, Mayo Clinic, Rochester, MN, US
130 Psychiatry and Behavioral Sciences, Stanford University School of Medicine, Stanford, CA, US
131 Rush University Medical Center, Chicago, IL, US
132 Scripps Translational Science Institute, La Jolla, CA, US
133 Neuroscience Research Australia, Sydney, NSW, AU
134 Faculty of Medicine, Department of Psychiatry, School of Health Sciences, University of Iceland, Reykjavik, IS
135 Division of Mental Health and Addiction, Oslo University Hospital, Oslo, NO
136 NORMENT, University of Oslo, Oslo, NO
137 Psychiatry and the Behavioral Sciences, University of Southern California, Los Angeles, CA, US
138 Mood Disorders, PsyQ, Rotterdam, NL
139 Institute for Medical Sciences, University of Aberdeen, Aberdeen, UK
140 Research Division, Federal Institute for Drugs and Medical Devices (BfArM), Bonn, DE
141 Centre for Addiction and Mental Health, Toronto, ON, CA
142 Neurogenomics, TGen, Los Angeles, AZ, US
143 Psychiatry, Psychiatrisches Zentrum Nordbaden, Wiesloch, DE
144 Computational Sciences Center of Emphasis, Pfizer Global Research and Development, Cambridge, MA, US
145 Department of Biostatistics, Princess Margaret Cancer Centre, Toronto, ON, CA
146 Dalla Lana School of Public Health, University of Toronto, Toronto, ON, CA
147 Psychological Medicine, Institute of Psychiatry, Psychology & Neuroscience, King's College London, London, GB
148 Department of Mental Health, Johns Hopkins University Bloomberg School of Public Health, Baltimore, MD, US
149 Institute of Genetic Medicine, Johns Hopkins University School of Medicine, Baltimore, MD, US
150 NORMENT, KG Jebsen Centre for Psychosis Research, Division of Mental Health and Addiction, Institute of Clinical Medicine and Diakonhjemmet Hospital, University of Oslo, Oslo, NO
151 National Institute of Mental Health, Klecany, CZ
152 Department of Psychiatry, University of Melbourne, Melbourne, Victoria, AU
153 Department of Psychiatry and Addiction Medicine, Assistance Publique - Hôpitaux de Paris, Paris, FR
154 Paris Bipolar and TRD Expert Centres, FondaMental Foundation, Paris, FR
155 UMR-S1144 Team 1: Biomarkers of relapse and therapeutic response in addiction and mood disorders, INSERM, Paris, FR
156 Psychiatry, Université Paris Diderot, Paris, FR
157 Psychiatry, University of Pennsylvania, Philadelphia, PA, US
158 Department of Psychiatry, University of Münster, Münster, DE
159 Division of Endocrinology, Children's Hospital Boston, Boston, MA, US
160 Centre for Affective Disorders, Institute of Psychiatry, Psychology and Neuroscience, London, GB
161 Department of Psychiatry & Psychology, Mayo Clinic, Rochester, MN, US
162 School of Medical Sciences, University of New South Wales, Sydney, NSW, AU

163 Department of Human Genetics, University of Chicago, Chicago, IL, US
164 Biometric Psychiatric Genetics Research Unit, Alexandru Obregia Clinical Psychiatric Hospital, Bucharest, RO
165 Institute of Neuroscience and Physiology, University of Gothenburg, Gothenburg, SE
166 INSERM, Paris, FR
167 Department of Medical & Molecular Genetics, King's College London, London, GB
168 Neuroscience Therapeutic Area, Janssen Research and Development, LLC, Titusville, NJ, US
169 Cancer Epidemiology and Prevention, M. Sklodowska-Curie Cancer Center and Institute of Oncology, Warsaw, PL
170 School of Psychology, The University of Queensland, Brisbane, QLD, AU
171 Research Institute, Lindner Center of HOPE, Mason, OH, US
172 Centre for Cognitive Ageing and Cognitive Epidemiology, University of Edinburgh, Edinburgh, GB
173 Human Genetics Branch, Intramural Research Program, National Institute of Mental Health, Bethesda, MD, US
174 Division of Mental Health and Addiction, Oslo University Hospital, Oslo, NO
175 Division of Mental Health and Addiction, University of Oslo, Institute of Clinical Medicine, Oslo, NO
176 Institute of Molecular and Cell Biology, University of Tartu, Tartu, EE
177 Mental Health, Faculty of Medicine and Health Sciences, Norwegian University of Science and Technology - NTNU, Trondheim, NO
178 Psychiatry, St Olavs University Hospital, Trondheim, NO
179 Psychosis Research Unit, Aarhus University Hospital, Risskov, DK
180 Munich Cluster for Systems Neurology (SyNergy), Munich, DE
181 University of Liverpool, Liverpool, GB
182 Psychiatry and Human Genetics, University of Pittsburgh, Pittsburgh, PA, US
183 Mental Health Services in the Capital Region of Denmark, Mental Health Center Copenhagen, University of Copenhagen, Copenhagen, DK
184 Division of Psychiatry, Haukeland Universitetssjukehus, Bergen, NO
185 Faculty of Medicine and Dentistry, University of Bergen, Bergen, NO
186 Human Genetics and Computational Biomedicine, Pfizer Global Research and Development, Groton, CT, US
187 College of Medicine Institute for Genomic Health, SUNY Downstate Medical Center College of Medicine, Brooklyn, NY, US
188 Department of Clinical Genetics, Amsterdam Neuroscience, Vrije Universiteit Medical Center, Amsterdam, NL
189 Department of Neurology and Neurosurgery, McGill University, Faculty of Medicine, Montreal, QC, CA
190 Montreal Neurological Institute and Hospital, Montreal, QC, CA
191 Department of Biomedical and NeuroMotor Sciences, University of Bologna, Bologna, IT
192 Department of Psychiatry, Massachusetts General Hospital, Boston, MA, US
193 Psychiatric and Neurodevelopmental Genetics Unit (PNGU), Massachusetts General Hospital, Boston, MA, US
194 Faculty of Medicine, University of Iceland, Reykjavik, IS
195 Department of Psychiatry, Hospital Namsos, Namsos, NO
196 Department of Neuroscience, Norges Teknisk Naturvitenskapelige Universitet Fakultet for naturvitenskap og teknologi, Trondheim, NO
197 Department of Genetics, University of North Carolina at Chapel Hill, Chapel Hill, NC, US
198 Department of Psychiatry, University of North Carolina at Chapel Hill, Chapel Hill, NC, US
199 Department of Psychiatry, McGill University, Montreal, QC, CA
200 Dept of Psychiatry, Sankt Olavs Hospital Universitetssykehuset i Trondheim, Trondheim, NO
201 Clinical Institute of Neuroscience, Hospital Clinic, University of Barcelona, IDIBAPS, CIBERSAM, Barcelona, ES
202 Institute of Biological Psychiatry, MHC Sct. Hans, Mental Health Services Copenhagen, Roskilde, DK
203 Department of Clinical Medicine, University of Copenhagen, Copenhagen, DK
204 Psychiatry, Indiana University School of Medicine, Indianapolis, IN, US
205 Biochemistry and Molecular Biology, Indiana University School of Medicine, Indianapolis, IN, US
206 Department of Pathology and Laboratory Medicine, University of California Los Angeles, Los Angeles, CA, US

### 1 Genome-wide association analyses in UK Biobank

To investigate the genetic architecture of psychiatric symptoms related to mood disturbance and psychosis, we used a novel combination of Bayesian item response theory and linear mixed models to conduct univariate GWASs in the UK Biobank ($N$ = 252,252). Details regarding the phenotypes and extensive quality control procedures are described below.

#### 1.1 Phenotype construction

Lifetime symptoms of depression and mania were assessed with two sets of items administered via in-person (Wave 1) and online (Wave 2) surveys. Lifetime symptoms of psychosis were only assessed during the online follow-up survey. Although items were very similar across assessment occasions, there were slight differences in wording and response options, as described below.

##### *1.1.1 Depression*

***Wave 1.*** The in-person surveys indexed depressive symptoms with two screener items that assessed the presence of notable, prolonged feelings of sadness or apathy.

1. "Looking back over your life, have you ever had a time when you were feeling depressed or down for at least a whole week?"
    - Data-Field 4598 (Data-Coding 100349).
    - Possible responses: "Yes", "No", "Do not know", and "Prefer not to answer".
2. "Have you ever had a time when you were uninterested in things or unable to enjoy the things you used to for at least a whole week?"
    - Data-Field 4631 (Data-Coding 100349).
    - Possible responses: "Yes", "No", "Do not know", and "Prefer not to answer".

***Wave 2.*** The web-based follow-up survey also indexed depressive symptoms with two screener items that assessed the presence of notable, prolonged feelings of sadness or apathy

1. "Have you ever had a time in your life when you felt sad, blue, or depressed for two weeks or more in a row?"
    - Data-Field 4598 (Data-Coding 503).
    - Possible responses: "Yes", "No", and "Prefer not to answer".
2. "Have you ever had a time in your life lasting two weeks or more when you lost interest in most things like hobbies, work, or activities that usually give you pleasure?"
    - Data-Field 4631 (Data-Coding 503).
    - Possible responses: "Yes", "No", and "Prefer not to answer".

##### *1.1.2 Mania*

***Wave 1.*** The in-person surveys indexed manic symptoms with two screener items that assessed the presence of notable, prolonged feelings of (hypo)mania or irritability.

1. "Have you ever had a period of time lasting at least two days when you were feeling so good, "high", excited or "hyper" that other people thought you were not your normal self or you were so "hyper" that you got into trouble?"
    - Data-Field 4642 (Data-Coding 100349).
    - Possible responses: "Yes", "No", "Do not know", and "Prefer not to answer".

2. "Have you ever had a period of time lasting at least two days when you were so irritable that you found yourself shouting at people or starting fights or arguments?"
    - Data-Field 4653 (Data-Coding 100349).
    - Possible responses: "Yes", "No", "Do not know", and "Prefer not to answer".

***Wave 2.*** The web-based follow-up survey indexed manic symptoms with two screener items that assessed the presence of notable, prolonged feelings of (hypo)mania or irritability.

1. "Have you ever had a period of time when you were feeling so good, "high", excited or "hyper" that other people thought you were not your normal self or you were so "hyper" that you got into trouble?"
    - Data-Field 20501 (Data-Coding 502).
    - Possible responses: "Yes", "No", "Do not know", and "Prefer not to answer".
2. "Have you ever had a period of time when you were so irritable that you found yourself shouting at people or starting fights or arguments?"
    - Data-Field 20502 (Data-Coding 502).
    - Possible responses: "Yes", "No", "Do not know", and "Prefer not to answer".

##### *1.1.3 Psychosis*

***Wave 2.*** The web-based follow-up survey assessed psychotic symptoms with four screener items. These items indexed various types of unusual beliefs and experiences that are indicative of psychosis or psychotic-like experiences.

1. "Did you ever see something that wasn't really there that other people could not see? Please do not include any times when you were dreaming or half-asleep or under the influence of alcohol or drugs."
    - Data-Field 20471 (Data-Coding 502).
    - Possible responses: "Yes", "No", "Do not know", and "Prefer not to answer".
2. "Did you ever hear things that other people said did not exist, like strange voices coming from inside your head talking to you or about you, or voices coming out of the air when there was no one around? Please do not include any times when you were dreaming or half-asleep or under the influence of alcohol or drugs."
    - Data-Field 20463 (Data-Coding 502).
    - Possible responses: "Yes", "No", "Do not know", and "Prefer not to answer".
3. "Did you ever believe that a strange force was trying to communicate directly with you by sending special signs or signals that you could understand but that no one else could understand (for example through the radio or television)? Please do not include any times when you were dreaming or half-asleep or under the influence of alcohol or drugs."
    - Data-Field 20474 (Data-Coding 502).
    - Possible responses: "Yes", "No", "Do not know", and "Prefer not to answer".
4. "Did you ever believe that there was an unjust plot going on to harm you or to have people follow you, and which your family and friends did not believe existed? Please do not include any times when you were dreaming or half-asleep or under the influence of alcohol or drugs."
    - Data-Field 20468 (Data-Coding 502).
    - Possible responses: "Yes", "No", "Do not know", and "Prefer not to answer".

##### *1.1.4 Item response theory models*

To construct psychiatric symptom phenotypes in UK Biobank, we used M*plus*[1] v8 to estimate a two-parameter probit multidimensional item response theory (IRT) model, which simultaneously combined self-report items across waves for each symptom dimension while leveraging all available information and accounting for the correlations between dimensions. IRT scaling was used as it does not assume that all items are equivalently related to the underlying construct of interest, accommodates differences in the base rate of item endorsement, and does not assume that missing data are missing completely at random. The advantages of IRT scaling using full-information estimation over averaging proportion scores for incomplete longitudinal data is described in more detail in Curran and colleagues[2]. The two parameter probit multidimensional IRT model can be expressed as:

$$P(y_{vij} = 1 | \theta_{vi}, \alpha_{vj}, \gamma_{vj}) = \Phi(\alpha_{vj}\theta_{vi} - \gamma_{vj}) = \int_{-\infty}^{\alpha_{vj}\theta_{vi} - \gamma_{vj}} \frac{1}{\sqrt{2\pi}} e^{\frac{-t^2}{2}} dt$$

where $P$ represents probability; $y_{vij}$ is the observed response for binary item $j$ for individual $i$ in symptom dimension $v$; and $\theta_{vi}$, $\alpha_{vj}$, and $\gamma_{vj}$ are scalar parameters that represent the latent construct hypothesized to underlie the observed item response patterns, item discrimination (*i.e.*, the degree to which the item discriminates between individuals in different regions on the latent continuum), and item difficulty (*i.e.*, the location where the item provides maximum information) for symptom dimension $v$. We freely estimated slope parameters for all items, and we modeled items as functions of single time-invariant, symptom-specific $\theta$ parameters.

We estimated IRT model parameters ($\alpha_{vj}$, $\gamma_{vj}$, and $\theta_{vi}$) using a Bayesian framework with non-informative priors and multiple imputation, which maximally leveraged all available responses for participants and minimized the impact of missing data. This approach is a full information approach that is asymptotically equivalent to maximum likelihood estimation. We imputed 100 plausible $\theta$ values for each participant to create a sampling distribution of $\theta$ for each symptom. The median values of these distributions were then used as the phenotype for the GWAS.

##### *1.1.5 Phenotypic factor structure in UK Biobank*

To evaluate the phenotypic factor structure of psychiatric symptoms and disorders characterized by mood disturbance and psychosis, we first created diagnostic phenotypes for the 252,252 individuals in the discovery sample, corresponding to diagnoses of major depressive disorder, bipolar disorder, schizoaffective disorder, and schizophrenia. These phenotypes were derived from electronic health records.

1. Diagnoses - main ICD-10
    - Data-Field 41202 (Data-Coding 19)
2. Diagnoses - secondary ICD-10
    - Data-Field 41204 (Data-Coding 19)

Due to low base rates, schizoaffective disorder and schizophrenia cases were combined. These variables were then included as observed variables in the multidimensional IRT model described in Supplementary Section 1.1.4. We first estimated the zero-order correlations between each pairwise combination of the psychiatric symptoms and disorders (Supplementary Figure 1). To maintain consistency across phenotypic and genetic factor analyses, we then conducted an exploratory factor analysis of the covariance matrix for these phenotypes, where we tested one-, two-, and three-factor solutions using the *factanal* function of R. As described in Supplementary Section 2.2, we retained the highest dimensional solution where each factor explained at least 20% of the variance.

Here, the three-factor solution satisfied this criterion (F1 = 35%, F2 = 29%, F3 = 23%), yielding three factors that were each predominantly characterized by a psychiatric disorder and their cardinal symptom(s). Specifically, we found that the factors corresponded to the following phenotypes (with loadings ≥ .30):

- F1: schizophrenia (1.08), psychosis (.90), and bipolar disorder (.31)
- F2: bipolar disorder (.54), mania (1.10), and depression (.51)
- F3: major depressive disorder (1.10) and depression (.31)

Collectively, these results suggest that symptoms and disorders tend to correlate most strongly in a canonical fashion at the phenotypic level (*i.e.*, psychosis with schizophrenia, mania with bipolar disorder, depression with major depressive disorder).

#### 1.2 Principal components of ancestry

Although the present analyses were limited to participants who self-reported non-Hispanic European descent, we sought to further account for cryptic relatedness and population stratification specific to this population. To this end, we used flashPCA2[3] to extract the first 40 principal components of ancestry in individuals who reported having non-Hispanic European ancestry.

We used an approach that broadly paralleled the original process used to estimate principal components of ancestry in the entire UKB dataset, as well as the default flashPCA2 recommendations as outlined by Abraham and colleagues in their online code repository (https://github.com/gabraham/flashpca). Specifically, we first generated a list of UKB participants that (i) were used in original PCA (*i.e.*, passed QC thresholds, pruned for kinship, etc.) and (ii) self-reported 'White British' ethnicity and have very similar genetic ancestry based on original PCA of the genotypes (as determined in the sample QC file provided by UKB investigators). We then extracted the hard genotype calls for those individuals and applied the recommended SNP-level QC thresholds (directly genotyped SNPs outside of long-range LD regions, minor allele frequency ≥ .01, genotyping call rate ≥ .02, missingness rate ≤ .05, and a Hardy-Weinberg equilibrium threshold ≥ 5e-6). Next, we applied the recommended LD pruning thresholds to produce a sample of 322,886 individuals with 77,355 independent markers before estimating the first 40 principal components with flashPCA2. Principal component loadings for each SNP used in the analysis were computed, exported, and then used to project all remaining participants of non-Hispanic European ancestry (e.g., siblings not used in the original PCA, participants of White Irish ancestry, participants of White Scottish ancestry, etc.) onto the PCs, yielding a final set of PC scores for the entire subsample. Scatterplots of the principal component scores were then manually examined to identify potential ancestral outliers in the sample.

#### 1.3 Quality control

We used an EasyQC[4] pipeline similar to the one reported by Linnér and colleagues[5] to check each set of UKB summary statistics for quality control problems. For each results file, we applied the following threshold in order.

1. We removed SNPs if either allele corresponded to a value other than “A”, “C”, “G”, or “T”.
2. We excluded SNPs if any of the following values were missing: *P* value, beta, standard error, effect allele frequency, sample size, and imputation accuracy (for imputed SNPs).
3. We excluded SNPs with values outside of permissible ranges (*e.g.*, negative or infinite standard errors, nonsensical *P* values, allele frequencies greater than 1 or below 0).
4. We dropped SNPs with minor allele frequencies less than .005.
5. We filtered out SNPs with low imputation accuracy, which was defined as an imputation score < .90.
6. We dropped duplicated SNPs based on GRCh37 base pair positions. When duplicated SNPs were identified, we retained the SNP with the largest sample size.

We then inspected several diagnostic plots to further ensure that results were not prone to systematic errors, including:

7. We checked for errors in allele frequencies and strand orientations by inspecting a plot of allele frequencies in our analytic sample against the allele frequency in a non-Hispanic European reference sample.
8. We checked for discrepancies between the reported $P$ values and the reported coefficient estimates and their SEs.
9. We looked for evidence of population stratification that had not been accounted for by checking a Q-Q plot.

### 2 Genomic structural equation modeling

Genomic structural equation modeling (Genomic SEM)[6] is a novel statistical method for applying structural equation modeling techniques to GWAS summary statistics to model the joint genetic architecture of complex traits. It is a flexible framework that allows for more accurate modeling of multivariate genetic covariance matrices, such as those derived from LD Score regression. Here, we conducted a series of Genomic SEM analyses to investigate the multivariate genetic architecture of the psychiatric symptoms and disorders characterized by mood disturbance and psychosis. The aim of these analyses is three-fold: (i) to identify the latent genetic factor(s) that best represent the factor structure of these phenotypes, (ii) to estimate the effects of individual SNPs on the latent genetic factor(s), and (iii) to evaluate heterogeneous effects among the discovery phenotypes.

#### 2.1 Hierarchical clustering

Hierarchical clustering is a form of cluster analysis that aims to identify features of a dataset that are similar to one another. It serves as a precursor to factor analysis or structural equation modeling, as its results can be used guide model specification decisions in subsequent analyses. To this end, we applied a hierarchical clustering algorithm to a genetic correlation matrix of our eight psychiatric phenotypes prior to any form of factor analysis. Specifically, we applied the complete-linkage hierarchical clustering algorithm employed by the *hclust* function of R. The algorithm identified two clusters present in the matrix.

The first cluster was comprised of the symptom-level phenotypes (depression, mania, and psychosis), major depressive disorder, and bipolar II disorder. The second cluster was comprised of bipolar I disorder, schizoaffective disorder, and schizophrenia. Beyond these two clusters, there clear evidence of a positive manifold across all items. All point estimates were positive, and all but two of the 28 pairwise genetic correlations were significant following Bonferroni correction.

#### 2.2 Factor analysis

Factor analysis is a multivariate statistical technique used to explain variance and covariance among sets of observed, correlated variables in terms of unobserved latent factors. It is a powerful tool for reducing dimensionality of data and accounting for measurement error in observed variables, often used in structural equation modeling. In factor analysis of genetic covariance matrices, $k$ observed variables are described as linear functions of $m$ latent variables, such that the model can be expressed as

$$y = \Lambda\eta + \varepsilon$$

where $y$ is a $k \times 1$ vector of observed variables, $\varepsilon$ is a $k \times 1$ vector of observed variable residuals, $\eta$ is a $m \times 1$ vector of latent variables, and $\Lambda$ is a $k \times m$ matrix of factor loadings that relate the observed variables to the latent variables.

In the present study, we used the *factanal* function of R to conduct an exploratory factor analysis with promax rotation. Guided by the results described in Supplementary Section 2.1, we tested factor solutions extracting up to three latent factors, retaining the highest dimensional solution where each factor explained at least 20% of the variance. As the three-factor solution did not meet this criterion (F1 = 43%, F2 = 26%, F3 = 18%), we selected the two-factor solution as the best exploratory factor model (Figure 2b).

In this two-factor solution, we found compelling evidence of approximate simple structure. Phenotypes principally loaded onto one of two latent genetic factors with negligible cross-loadings. The two correlated latent genetic factors explained 81.3% of the total genetic variance across phenotypes.

#### 2.3 Structural equation modeling

Structural equation modeling is a statistical framework comprised of a diverse set of models and methods used to explain variance and covariance among sets of variables. While the background and many applications of structural equation modeling are extensive[7,8], the fundamentals as they relate to the Genomic SEM framework are briefly reviewed below.

Structural equation models can be represented in two sets of equations: the *measurement model*, which describes how observed variables relate to latent variables, and the *structural model*, which describes how latent variables relate to one another[6]. As in exploratory factor analysis, $k$ observed variables are again described as linear functions of $m$ continuous latent variables. In confirmatory factor analysis, this is referred to as the measurement model, which is still expressed as

$$y = \Lambda\eta + \varepsilon$$

with the same notation as described in Supplementary Section 2.2.

If theory is used to explain associations between latent variables, a structural model can then be specified to relate latent variables to each other via directed regression coefficients. The structural model can be expressed as

$$\eta = \mathrm{B}\eta + \zeta$$

where B is a $m \times m$ matrix of regression coefficients that relate latent variables to one another and $\zeta$ is a $m \times 1$ vector of latent variable residuals. In this full structural equation model, the observed sample covariance matrix is represented by a set of parameters that relates observed variables to latent variables, and latent variables to each other in a series of linear equations.

Genomic SEM employs a two-stage SEM approach to model the genetic covariances between a set of phenotypes (*i.e.*, the observed phenotypes). Stage 1 consists of estimating the genetic covariance matrix and the sampling covariance matrix. Stage 2 consists of fitting a structural equation model that minimizes misfit between the model-implied and empirical genetic covariances.

In Stage 1, Genomic SEM uses a multivariable form of LD Score regression to estimate the empirical genetic covariance matrix ($S$) and its associated sampling covariance matrix ($V_S$). Here, $S$ is a symmetric matrix of order $k$ with SNP heritabilities on the diagonal and genetic covariances between phenotypes off the diagonal. Comprised of $k^* = \frac{k(k+1)}{2}$ nonredundant elements, $S$ can be written as

$$S = \begin{bmatrix} h_1^2 & & & \\ \sigma_{g1.g2} & h_2^2 & & \\ \vdots & & \ddots & \\ \sigma_{g1,gk} & \sigma_{g2,gk} & \dots & h_k^2 \end{bmatrix}$$

To obtain unbiased estimates of standard errors and test statistics, Genomic SEM then constructs the asymptotic sampling covariance matrix of the LD Score regression estimates, $V_S$, by using all nonredundant elements in the $S$ matrix. Here, $V_S$ is a symmetric matrix of order $k^*$where the diagonal elements are sampling variances and the off-diagonal elements are sampling covariances. Thus, it can be written as

$$V_S = \begin{bmatrix} SE(h_1^2)^2 & & & & & \\ cov(h_1^2, \sigma_{g_1,g_2}) & SE(\sigma_{g_1,g_2})^2 & & & & \\ \vdots & \vdots & \ddots & & & \\ cov(h_1^2, \sigma_{g_1,g_k}) & cov(\sigma_{g_1,g_2}, \sigma_{g_1,g_k}) & SE(\sigma_{g_1,g_k})^2 & & & \\ \vdots & \vdots & \vdots & \ddots & & \\ cov(h_1^2, h_j^2) & cov(\sigma_{g_1,g_2}, h_j^2) & cov(\sigma_{g_1,g_k}, h_j^2) & SE(h_j^2)^2 & & \\ \vdots & \vdots & \vdots & & \ddots & \\ cov(h_1^2, \sigma_{g_j,g_k}) & cov(\sigma_{g_1,g_2}, \sigma_{g_j,g_k}) & cov(\sigma_{g_1,g_k}, \sigma_{g_j,g_k}) & cov(h_j^2, \sigma_{g_j,g_k}) & SE(\sigma_{g_j,g_k})^2 & \\ cov(h_1^2, h_k^2) & cov(\sigma_{g_1,g_2}, h_k^2) & cov(\sigma_{g_1,g_k}, h_k^2) & cov(h_j^2, h_k^2) & cov(\sigma_{g_j,g_k}, h_k^2) & SE(h_k^2)^2 \end{bmatrix}$$

The diagonal elements of $V_S$ are then estimated with a jackknife resampling procedure following the original bivariate version of LD Score regression.

In Stage 2, Genomic SEM uses the $S$ and $V$ matrices from Stage 1 to estimate the parameters of the specified structural equation model using a weighted least squares (WLS) fit function, as detailed in Grotzinger et al. (2019). Notably, this method is capable of accounting for differences in GWAS sample size, which is ideal in the present study. Furthermore, the off-diagonal elements of $V_S$ index the extent to which the sampling errors across input GWAS are correlated, which means that Genomic SEM, like LD score regression upon which it is based, is unbiased and robust to varying degrees of, or even complete, sample overlap.

##### *2.3.1 Confirmatory factor analysis*

Confirmatory factor analysis is a common application of structural modeling where theoretical models are used to explain the observed covariances among a set of observed variables. Here, we tested a series of competing models to identify the model that best fit the data, where good fit indicated that the specified latent variable structure adequately explained the observed genetic covariances among the set of observed variables.

Guided by the results described in Supplementary Sections 2.1 and 2.2, as well as psychiatric and psychometric theory, we tested a series of confirmatory factor models to identify the solution that best explained the observed genetic covariances among the set of discovery phenotypes. Specifically, we tested four models: (i) a single common factor model (*i.e.*, a *p* factor), (ii) a correlated factors model (iii) a correlated factors model with bipolar II disorder to cross-loading on both factors, and (iv) a correlated factors model with correlated residuals between bipolar I disorder and bipolar II disorder. Path diagrams for these models are presented in Supplementary Figure 2. Unit variance identification was used to set the scale of the latent factors.

Model fit was assessed using conventional indices in structural equation modeling: the model $\chi^2$ statistic, the Akaike information criterion (AIC), the comparative fit index (CFI), and the standardized root mean square residual (SRMR). All of these indices retain their standard interpretations within a Genomic SEM framework with the exception of the model $\chi^2$ statistic [6]. In large samples, such as those used here, $\chi^2$ tests are overpowered and likely to be significant. As such, the model $\chi^2$ statistic was used as a comparative measure of fit to evaluate competing models (akin to AIC), rather than a measure of statistical significance. For CFI and SRMR, values greater than .90 and less than .08, respectively, were considered reflective of good model fit[9].

***Common factor model***
As a baseline, we evaluated a common factor model with all eight phenotypes operating as indicators for a single latent factor. While easily interpretable, this particular model exhibited poor fit, as indicated by model fit indices ($\chi^2(20)$ = 1630.24, AIC = 1662.24, CFI = .99, SRMR = .20). Inspection of the observed and model implied genetic correlation matrices indicated that the common factor model implied that genetic correlations between schizophrenia, schizoaffective disorder, bipolar I disorder, and bipolar II disorder were severely downwardly biased. Moreover, the genetic correlations between psychiatric symptoms and disorders were modestly upwardly biased in the common factor model.

***Correlated factors model***
Preliminary results described in Supplementary Sections 2.1 and 2.2 suggested the promise of a two-factor model, where phenotypes loaded onto two distinct-but-correlated latent genetic factors. As an initial test of this model, we estimated a simple correlated factors model with no cross-loadings and no correlated residuals. While this model had better fit than the common factor model, some model fit indices were still suboptimal ($\chi^2(19)$ = 608.97, AIC = 642.97, CFI = .99, SRMR = .11). We next fit a model allowing bipolar II disorder to cross-load onto both factors. This model also showed good fit, as indicated by model fit indices ($\chi2(18)$ = 390.93, AIC = 426.93, CFI = .99, SRMR = .08), but the cross-loading resulted in markedly lower loadings for bipolar II disorder (.51 and .43 for F1 and F2, specifically). Finally, we fit a correlated factors model that estimated the correlation between residual variance in bipolar I disorder and bipolar II disorder. This model fit the data best, closely approximating the observed genetic covariance matrix ($\chi2(18)$ = 496.16, AIC = 532.16, CFI = .99, SRMR = .06).

While all variations of the correlated factors model showed improved fit over the common factor model, we identified the model that included correlated residuals as the best fitting model. Notably, this model provided a parsimonious and easily interpretable factor structure that simultaneously minimized the standardized difference between the observed and predicted genetic correlations. Inspection of the observed and model implied genetic correlation matrices indicated that the final correlated factors model fit the data substantially better than the common factor model because it appropriately accounted for different patterns of covariance by segregating the phenotypes into two separate-but-correlated latent factors (Supplementary Figure 3).

##### *2.3.2 Genetic correlation*

Structural equation models can also be used to estimate the genetic correlation between an unobserved latent factor and an observed exogenous phenotype not included in the model. Indeed, this method is preferable to using bivariate LD score regression, as it based on the genetic covariances directly rather than the estimated SNP effects, which may not be mediated by the latent factor(s) (see Supplementary Section 2.5 for more on heterogenous SNP effects). To estimate genetic correlations between a latent factor and an observed exogenous phenotype, we first created a single-item quasi-latent factor for the exogenous phenotype, and fixed the residual variance for the phenotype to zero. We then estimated the correlation between our latent factors and the quasi-latent factor for the phenotype of interest.

##### *2.3.3 Multivariable genetic regression*

The approach described in Supplementary Section 2.3.2 can be extended to conduct multivariable genetic regression, which yields estimates of genetic associations between two variables after accounting for relationships with additional variables in the model (*i.e.*, partial genetic correlations). Here, this is done by regressing the exogenous phenotype onto F1 and F2 while simultaneously estimating the genetic correlation between the two latent genetic factors.

#### 2.4 Multivariate genome-wide association analyses

After identifying a confirmatory factor model that best explained the observed genetic covariances among the phenotypes, we conducted a multivariate GWAS by estimating the individual SNP effects on each latent factor in the model. A brief overview of this method is provided below.

To estimate the effect of SNP $j$ on F1 and F2, individual SNP effects are included in both the genetic covariance matrix and the sampling covariance matrix. This is accomplished by expanding the genetic covariance matrix to include covariances between SNP $j$ and the latent genetic components of each phenotype, $g_1$ through $g_k$.

$$S = \begin{bmatrix} \sigma^2_{SNP} & & & \\ \sigma_{SNP,g_1} & h^2_1 & & \\ \sigma_{SNP,g_2} & \sigma_{g_1,g_2} & h^2_2 & \\ \sigma_{SNP,g_k} & \sigma_{g_1,g_k} & \sigma_{g_2,g_k} & h^2_k \end{bmatrix}$$

The associated sampling covariance matrix, $V_S$, then includes the following: (i) the sampling variances and sampling covariances of the SNP heritabilities and genetic covariances, (ii) the variance of SNP $j$ as derived from reference panel data, and (iii) the sampling covariances of the SNP-genotype covariances. Finally, Genomic SEM is used to estimate $m$ models to obtain GWAS summary statistic for the latent factors, where $m$ is the number of SNPs present across all included summary statistics.

Note that unit loading identification is used to set the scale of latent factors for models including SNP effects. This is a difference from the structural equation models without SNP effects, where unit variance identification is used to facilitate easy interpretation of factor loadings. This is done for two reasons. First, if the variance of the factor were set to 1, the inclusion of a SNP as a regressor technically changes the variance of the latent factor to be 1 plus the variance explained by the SNP. Second, the use of unit variance identification scales SNP effects as if they were for a phenotype that was entirely heritable (*i.e.*, $h^2_{SNP} = 1$). This distinction does not change the ratio of effect estimates to standard errors, but it does potentially complicate comparison to other GWAS results. Thus, we find that unit loading identification is the most appropriate method to scale latent factors in models that include SNP effects. Here, we set the scale of F1 and F2 by fixing the factor loadings of psychotic symptoms and schizophrenia, respectively.

#### 2.5 Heterogeneity tests

It is possible that SNP effects might vary across each indicator and not act entirely through the common latent variable. To evaluate this potential heterogeneity in SNP effects, we computed genome-wide $Q_{\text{SNP}}$ statistics, which are $\chi^2$-distributed test statistics estimated for each SNP in the multivariate GWAS. As described by Grotzinger and colleagues[6], larger values for $Q_{\text{SNP}}$ reflect a violation of the null hypothesis that the SNP acts entirely through the latent factor(s).

#### 2.6 Effective sample size

While it can be difficult to estimate effective sample size for a given SNP in a latent factor model, we were able to produce reasonable estimates of effective sample size for the overall multivariate GWAS under a set of reasonable assumptions. First, we assume that the effect of SNP $j$ follows

$$\beta_j = \frac{Z_j}{\sqrt{n_j \times 2 \times MAF_j \left(1 - MAF_j\right)}}$$

where $Z$ is the $Z$ statistic, $n$ is the unknown effective sample size that we seek to estimate, and $MAF$ is the minor allele frequency of SNP $j$. Note that the variance of SNP $j$ ($\sigma_j^2$) is estimated as $2 \times MAF_j \left(1 - MAF_j\right)$. Therefore, if we know the effect and MAF of that SNP, then we can estimate its effective sample size by solving for $n_j$.

$$\frac{\beta_j}{Z_j} = \frac{1}{\sqrt{n_j \times \sigma_j^2}}$$

$$\frac{Z_j}{\beta_j} = \sqrt{n_j \times \sigma_j^2}$$

$$\left(\frac{Z_j}{\beta_j}\right)^2 = n_j \times \sigma_j^2$$

$$n_j = \frac{(Z_j/\beta_j)^2}{\sigma_j^2}$$

This formula will typically produce reasonable estimates of $n_j$, but it can be prone to error for SNPs with low MAF. Here, we set a lower and upper MAF limit of approximately 10% and 40%, respectively, when estimating effective N for the overall multivariate GWAS results ($N_{eff}$). Following this, we estimate that $N_{eff}$ is approximately equal to the mean $n_j$ for $m$ SNPs with a MAF between $a$ and $b$. This can be expressed as

$$N_{eff} \approx \frac{1}{m} \sum_{MAF=a}^{b} n_j$$

Here, we apply this to the results for the results for F1 and F2 and estimate that the effective $N$ for each phenotype is 377,518 and 51,276, respectively. We note that this calculation is robust to sample overlap in the multivariate GWAS, as Genomic SEM accounts and corrects for such overlap.

We note that with an effective sample size, it is possible to backout an estimate of genetic variance that is, in one sense, conceptually analogous to SNP heritability in that they reflect the scale of SNP effects for a GWAS target. But there is no information about phenotypic variance of the latent genetic factors, and so should not be interpreted as heritabilities. Rather, these genetic variance estimates are most useful as an additional metric for comparing GWAS results – especially when paired with effective sample size. Here, the latent factors F1 and F2 have genetic variance estimates of 6% (SE = .28%) and 56% (SE = 2.29%) respectively. The genetic variance estimates for F1 and F2 differ because of the differing SNP heritabilities of their underlying indicators, with lower heritabilities observed for self-report symptoms than for clinically-defined disorders. The greater proportion of non-genetic variance in self-report symptoms might be due to greater influence of environmental variation and/or greater unreliability of measurement.

### 3 Supplementary figures

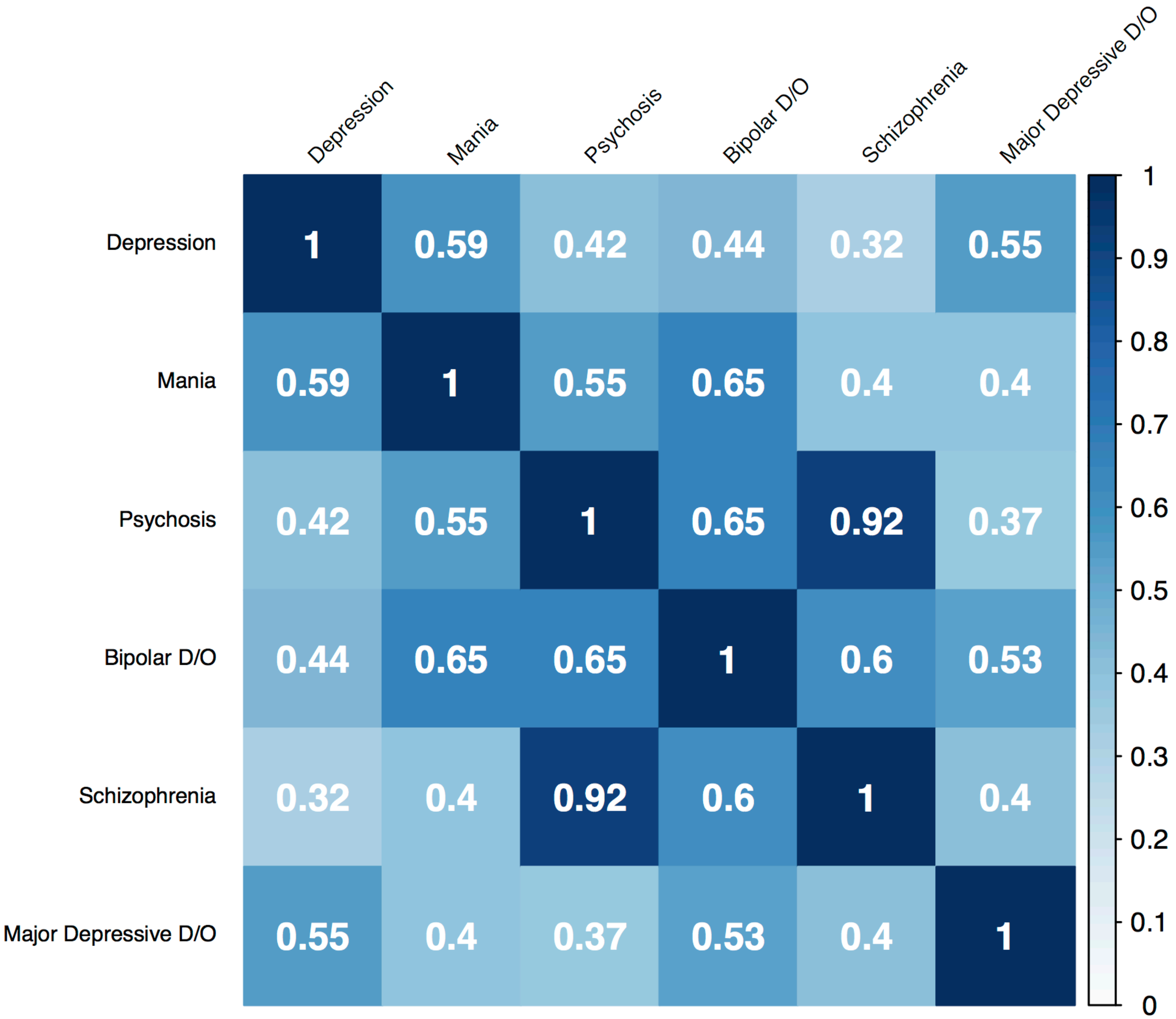

**Supplementary Figure 1. Phenotypic correlations between psychiatric symptoms and disorders.** Matrix of phenotypic correlations for available psychiatric symptoms and disorders in the UK Biobank discovery sample.

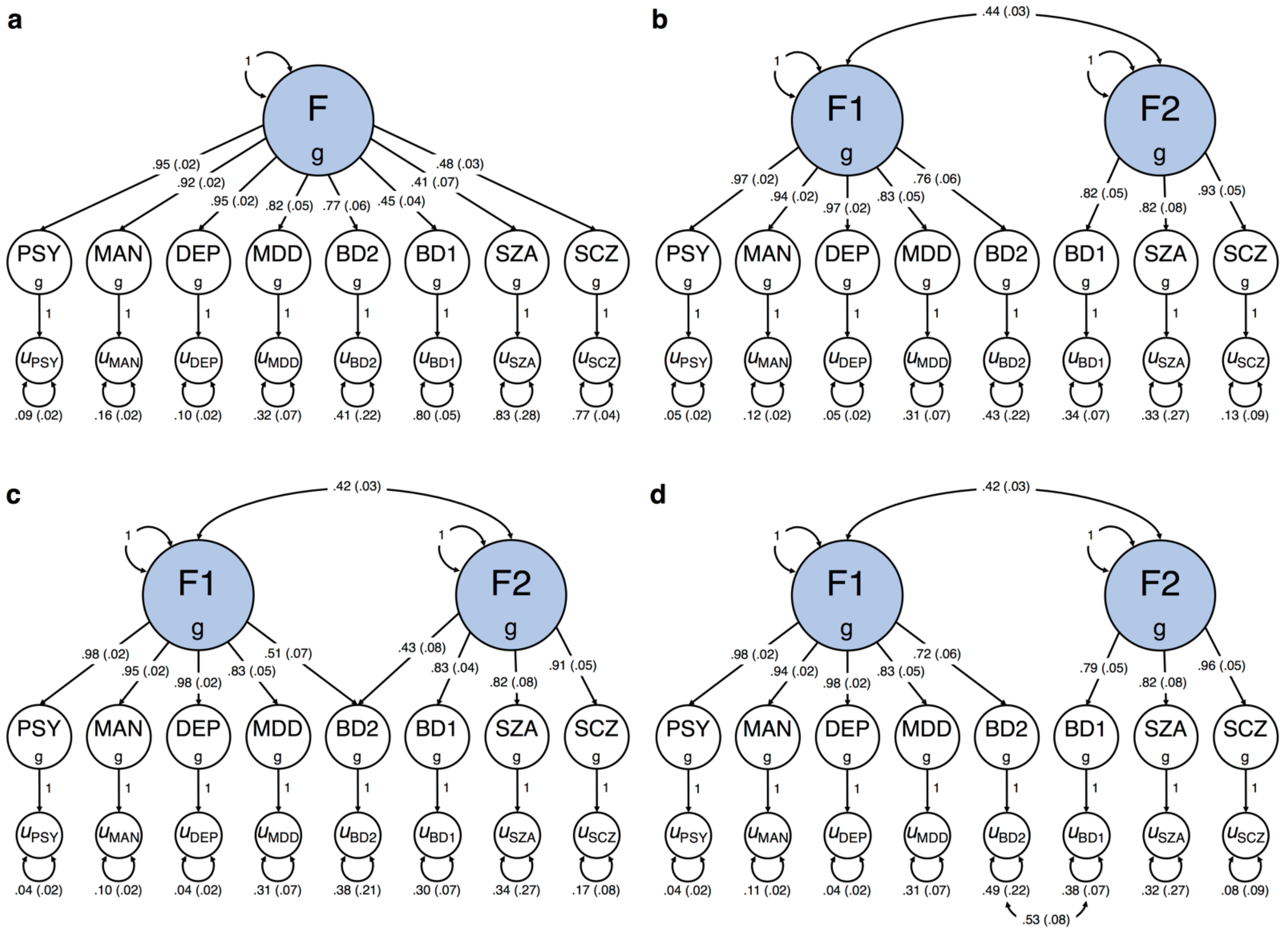

**Supplementary Figure 2. Path diagrams for the four confirmatory factor models. a**,**b**,**c**,**d,** Path diagrams with standardized parameter estimates for (**a**) a common factor model, (**b**) a correlated factors model, (**c**) a correlated factors model with bipolar II disorder cross-loading on both factors, and (**d**) a correlated factors model with correlated residuals between bipolar I disorder and bipolar II disorder. Standard errors are reported in parentheses next to each parameter estimate. Latent genetic factors are highlighted in blue.

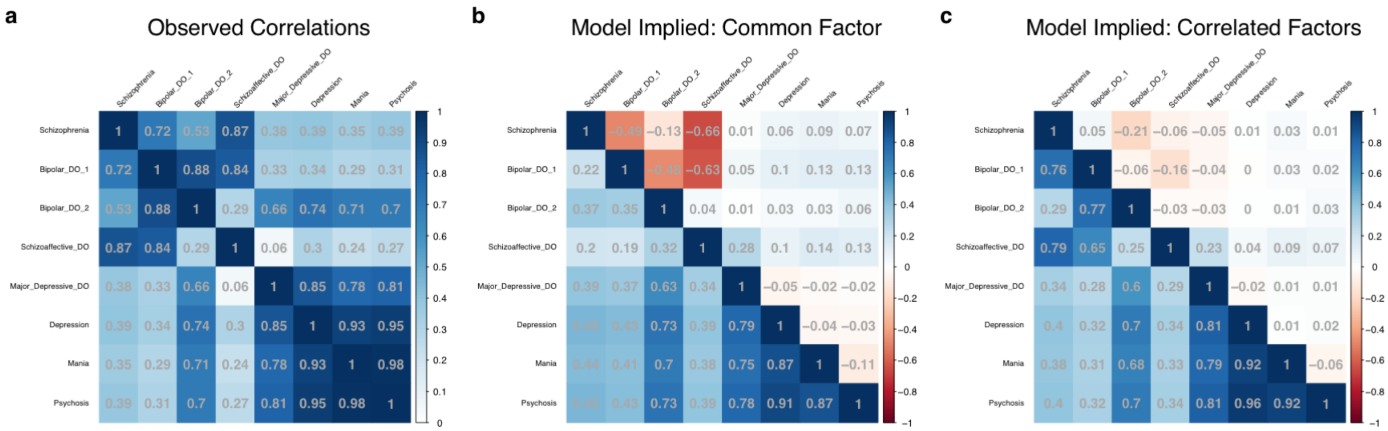

**Supplementary Figure 3. Observed and implied genetic correlation matrices. a**, Matrix of observed genetic correlations for the eight psychiatric symptoms and disorders. **b**,**c**, Matrices of model implied genetic correlations for (**a**) the common factor (*i.e.*, the *p* factor) and (**b**) the final correlated factors model (Supplementary Figure 2d). Model implied correlations are presented below the diagonal while the difference between the model implied and observed correlations (model implied $r_g$ – observed $r_g$) are presented above the diagonal. Positive and negative values above the diagonal, therefore, reflect upwardly and downwardly biased estimates, respectively.

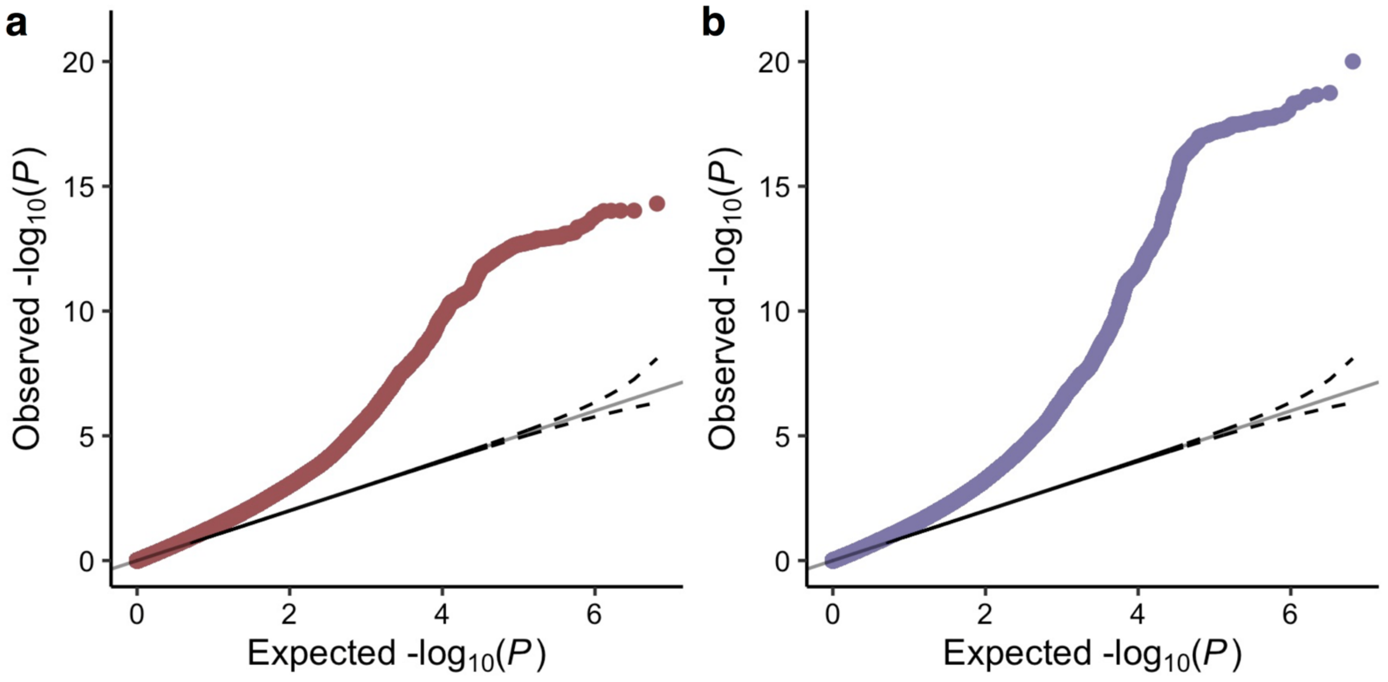

**Supplementary Figure 4. Quantile-quantile plot for the latent genetic factors. a**,**b**, Quantile-quantile plot for (**a**) F1 and (**b**) F2, illustrating strong polygenic signal for both multivariate GWAS. The y-axis corresponds to the observed distribution of P, while the x-axis corresponds to the expected distribution of P under the null. The null is plotted as a solid gray line, and the accompanying 95% confidence interval is plotted as a dotted black line.

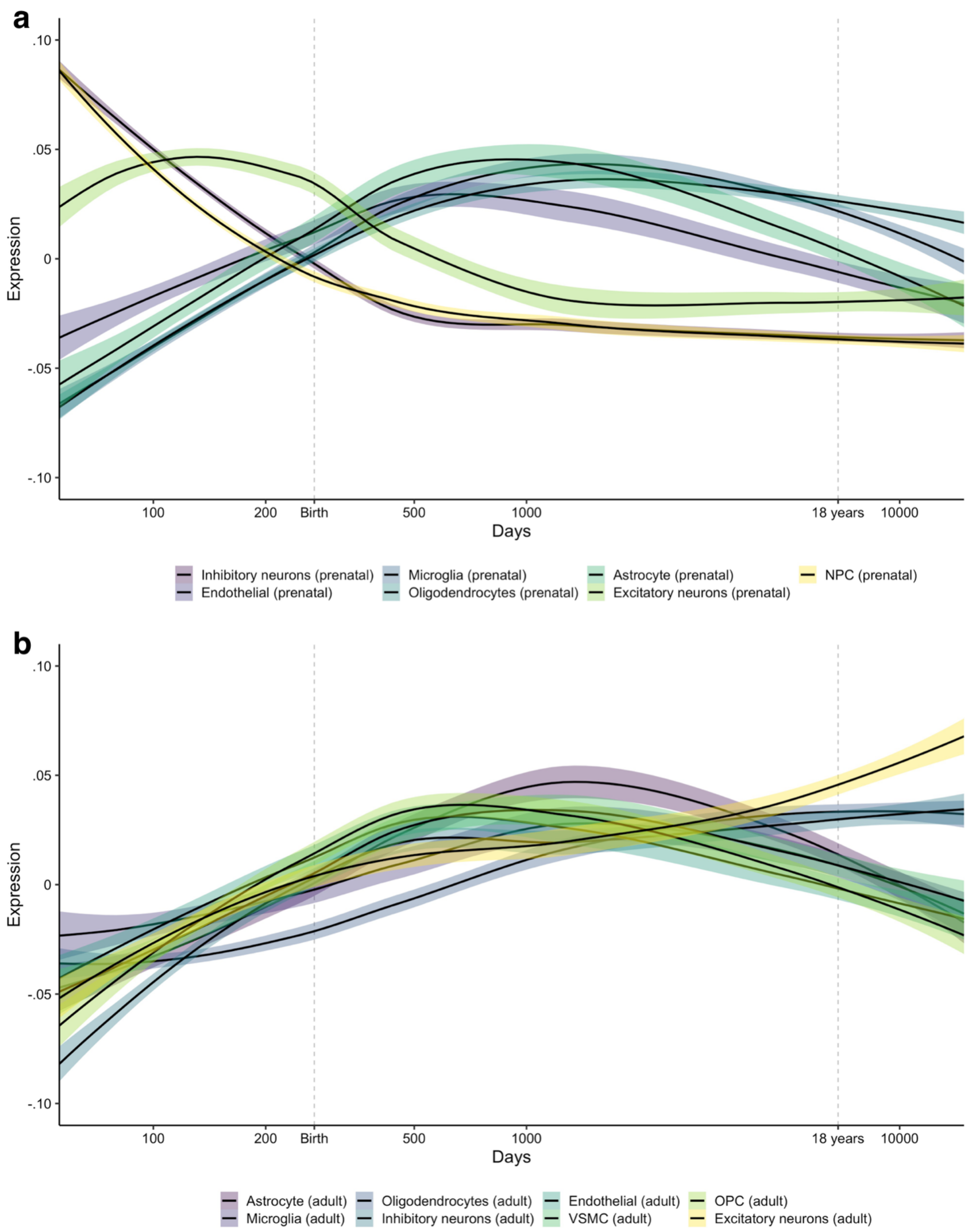

**Supplementary Figure 5. Spatiotemporal gene expression of specific cell types. a**,**b**, Developmental trajectories fit via LOESS regression for (**a**) prenatal and (**b**) adult cell types in the Brainspan dataset. Cell type data is derived from Li and colleagues[10]. Note: inhibitory neurons = mostly GABAergic interneurons, excitatory neurons = mostly glutamatergic excitatory projection neurons, NPC = neural progenitor cells, VSMC = vascular smooth muscle cells, OPC = oligodendrocyte progenitor cells.
